# Rapid robust high-fidelity 3D neuronal extraction from multi-view projections

**DOI:** 10.64898/2025.12.01.691740

**Authors:** Yujia Chen, Guoxun Zhang, Mingrui Wang, Yuanlong Zhang, Jingyu Xie, Zhifeng Zhao, Ruqi Huang, Jiamin Wu, Qionghai Dai

## Abstract

Recent developments in imaging facilitate large-scale three-dimensional (3D) neuronal recording. While these huge amounts of data shed new light on population-level neural coding, it is much harder to extract neuronal calcium dynamics from 3D volumes than 2D images with the existence of noise and scattering. Here, we presented DeepWonder3D, a general end-to-end pipeline for rapid and robust 3D neuronal extraction with high fidelity. Instead of processing voxel by voxel, DeepWonder3D works on the multi-view projections of 3D imaging data obtained either digitally or optically through specific point spread functions (PSFs) for general applicability to diverse techniques, including point-scanning microscopy, light-field microscopy, and two-photon synthetic aperture microscopy. Integrating denoising, resolution registration, background removal, neuronal extraction, and multi-view fusion into a unified pipeline tailored for large-scale high-resolution datasets contaminated by noise and scattering, DeepWonder3D outperforms state-of-the-art methods in 3D localization accuracy with a 10-fold reduction in computational costs, validated by numerical simulations and a hybrid two-photon/light-field imaging system. With the RUSH3D mesoscope, DeepWonder3D achieves rapid high-fidelity 3D calcium extraction of tens of thousands of neurons across the mouse cortex within hours.

## Introduction

Understanding the relationship between neural circuit activity and complex functions such as perception, cognition, and behavior is a central goal in neuroscience^1,2^. The mammalian brain comprises billions of neurons forming intricate networks with trillions of synaptic connections, and its functions arise from the dynamic patterns of electrical and chemical signaling within these circuits^3^. Elucidating these patterns is fundamental not only for basic science but also holds important relevance for understanding the pathophysiology of neurological and psychiatric disorders. A key requirement for advancing this understanding is the ability to monitor the activity of large, spatially distributed neuronal populations *in vivo* with high spatial and temporal resolution^4–6^.

Fluorescence microscopy with genetically encoded calcium indicators (e.g., GCaMP) enables less invasive, large-scale optical monitoring of calcium dynamics at single-cell resolution within genetically targeted populations in living organisms^7^. However, high-speed three-dimensional (3D) calcium imaging is essential for studying distributed neural circuits, as neurons are embedded in complex 3D structures *in vivo*^8^. Fortunately, recent advances in computational imaging have enabled 3D neuronal recording by engineering the PSF, allowing imaging of neuronal activity throughout the 3D structure of the cortex with high spatial and temporal resolution. Mathematically, a bounded 3D volume can be implicitly represented as a set of all multi-view projections 𝓥 = {V(u,v)(x,y)}(u,v) ∈ Ω, where each V(u,v)(x,y) is a two-dimensional (2D) projection of the 3D volume from the viewpoint (u,v), and (x,y) denotes the spatial coordinates on the image plane^9^. Hence, many techniques encode 3D spatial information into multiple views captured on a 2D sensor, from which the full 3D volume can be computationally reconstructed. For instance, light field microscopy (LFM)^10^ employs a microlens array to acquire snapshot volumetric images, offering higher volumetric frame rates than conventional point- or plane-scanning methods^11^. Two-photon synthetic aperture microscopy (2pSAM)^12^ adapts the synthetic aperture principle from radar systems to two-photon microscopy, enabling millisecond-scale 3D imaging of subcellular activity in deep tissue across more than 100,000 volumes. RUSH3D mesoscope^13^ integrates digital adaptive optics (DAO) to support rapid, large-volume imaging, capturing dynamics across expansive fields of view (e.g., 8,000×6,000×400 µm³at 20 Hz) with high throughput and reduced system complexity. These advanced 3D calcium imaging techniques have facilitated various high spatiotemporal resolution *in vivo* applications, such as large-scale neuronal calcium imaging^12–14^, dynamic imaging of zebrafish hearts^15–17^, voltage imaging of individual neurons^18^, and 3D single-cell analysis^19^.

Multi-view projections offer an attractive strategy for rapid volumetric imaging of neuronal activity, providing high temporal resolution across extended volumes^20^. However, performing 3D neuronal extraction on multi-view projections of 3D calcium imaging data is not easy. Unlike 2D methods, 3D neuronal extraction requires reconstructing the 3D cortex from multiple 2D views. While volume reconstruction is the theoretically ideal way to analyze multi-view projections and restore its full volumetric information, practical application in dense tissues like the mammalian cortex faces significant hurdles^21^. The primary obstacle is photon scattering, which creates pervasive background haze, reduces spatial resolution and contrast, and severely undermines the accuracy of 3D neuronal localization^22^. Reconstructing a clear 3D volume from this scattered light is considerably difficult. First, this task is an ill-posed inverse problem^21^, meaning a stable solution is theoretically challenging to achieve. Using conventional deconvolution approaches to solve the ill-posed inverse problem tends to amplify noise or introduce new reconstruction artifacts^23^. Second, these methods typically fail to fully leverage the rich information in multi-view projections^24^, especially with the low signal-to-noise ratios common in deep-tissue imaging. Consequently, relying on these flawed, pre-reconstructed 3D volumes for neuronal extraction only further compounds these inaccuracies. Moreover, high-resolution 3D calcium imaging inherently generates extremely large data of multi-view recordings, particularly during long-term or large-volume acquisitions, imposing substantial burdens on data storage^25^ and computational processing^26^. These intertwined challenges of image degradation from noise and scattering, reconstruction inaccuracies, and heavy data and computational loads severely pose substantial obstacles to 3D neuronal extraction from multi-view projections. Therefore, there is an urgent need for a more advanced and computationally efficient 3D neuronal extraction workflow to enable high-fidelity, large-scale mapping of 3D neuronal populations in deep, optically scattering tissues using multi-view projections of 3D calcium imaging data.

We developed DeepWonder3D, an integrated framework based on self-supervised learning^27^ and simulation-supervised learning^28^, designed for the rapid and robust 3D neuronal extraction from multi-view projections of 3D calcium imaging data (Fig. 1a). To improve the robustness of our method, we recognized the need to tackle scattering and noise upfront, as these factors can negatively interfere with accurate neuronal extraction. Therefore, instead of processing voxel by voxel, DeepWonder3D operates directly on multi-view projections and sequentially performs denoising, resolution registration, background removal, neuronal extraction, and multi-view fusion. By applying essential preprocessing steps directly to the multi-view projections prior to 3D neuronal extraction, DeepWonder3D effectively mitigates the detrimental impacts of noise and scattering. As these preprocessing steps lead to greater consistency in the quality of recordings, this strategy not only enhances the robustness of neuronal extraction but also confers generalizability to DeepWonder3D across diverse microscopy systems and various imaging conditions (Fig. 1b, 1c). Operating directly on the multi-view projections allows DeepWonder3D to fully leverage the rich information within the 3D calcium imaging data while circumventing potential artifacts often introduced during conventional volume reconstruction, thereby improving the accuracy of 3D neuronal localization during our multi-view fusion. Since data from the 3D cortex can be effectively projected into multi-view representations, DeepWonder3D offers a general solution for 3D neuronal extraction from different kinds of 3D neuronal recordings. Furthermore, to achieve a rapid workflow essential for large-scale neurophysiological studies, DeepWonder3D integrates deep learning to streamline the analytical workflow and leverages TensorRT to further accelerate neuronal extraction. To rigorously assess the performance of DeepWonder3D, we conducted quantitative evaluations on both simulated and experimental datasets, measuring 3D localization accuracy, temporal correlation, and computational efficiency. For experimental validation, we developed a hybrid imaging platform integrating single-photon (1p) LFM with two-photon (2p) microscopy. Furthermore, the scalability and generalizability of DeepWonder3D to diverse imaging modalities was demonstrated through successful application to data acquired using 2pSAM and RUSH3D mesoscope, confirming its utility for multi-modal analysis.

**Figure 1.**
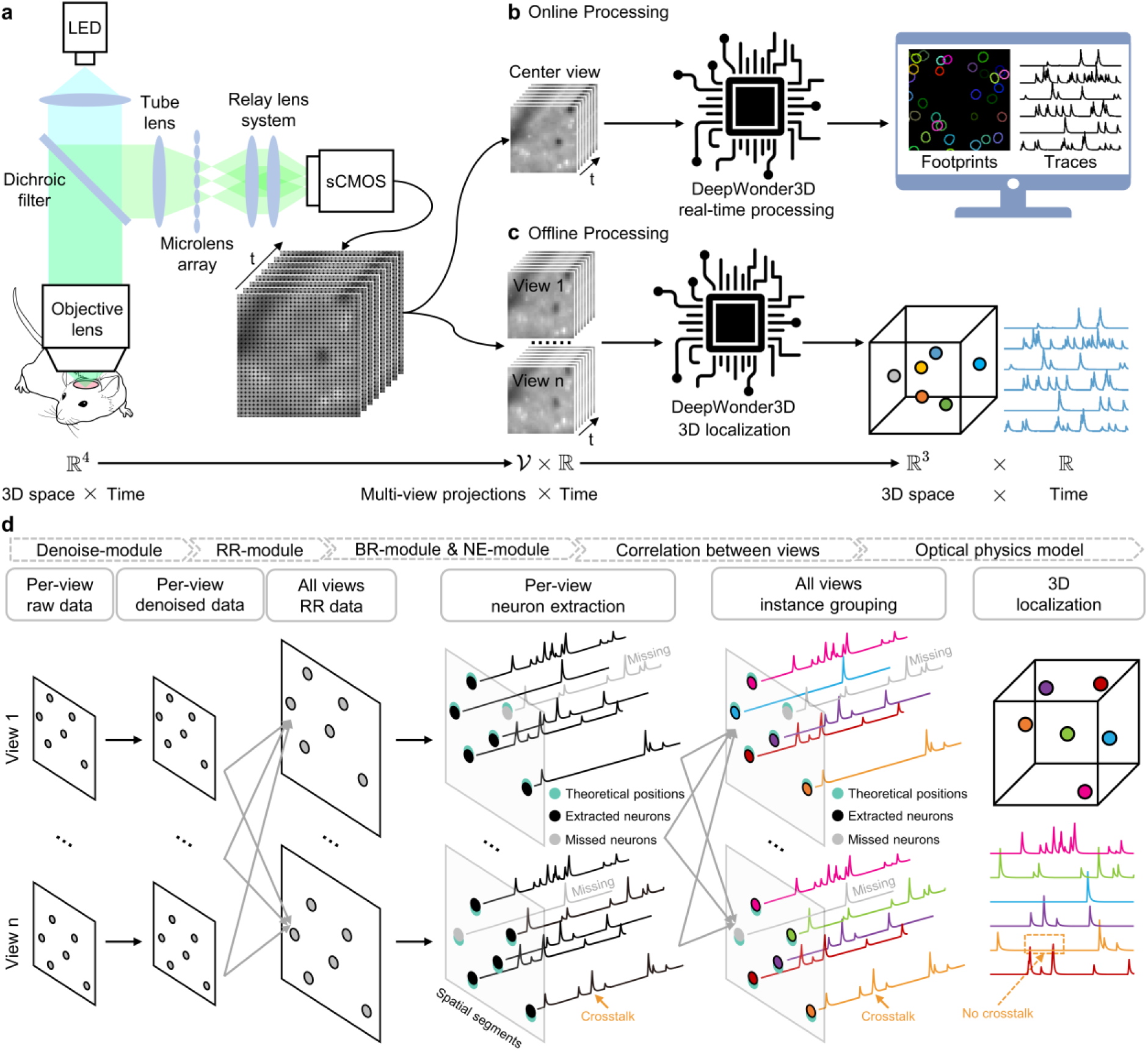
Principle and workflow of DeepWonder3D. a, Schematic illustration of the light field imaging system. The system comprises an LED light source, a dichroic filter, a tube lens, a relay lens system, a microlens array, and an objective lens. This configuration enables the capture of volumetric fluorescence signals, indicative of neural activity, onto a camera. The imaging system performs a mapping from the 3D space- time domain (ℝ^4^) to the multi-view projection-time domain (𝓥 × ℝ). b, Workflow for online processing and real-time analysis. The DeepWonder3D workflow processes center-view images to extract neuronal spatial locations and corresponding temporal activity traces for immediate analysis. c, Workflow for offline processing and comprehensive 3D localization. DeepWonder3D processes multi-view image stacks to accurately reconstruct neuronal positions in 3D space (ℝ^3^) and extract their activity traces over time (ℝ). DeepWonder3D maps the multi-view projection-time domain (𝓥 × ℝ) back to the 3D space-time domain (ℝ^4^), effectively also disentangling the inherent mixed patterns within the original ℝ^4^ domain by factorizing them into spatial (ℝ^3^) and temporal (ℝ) representations. d, Stepwise processing workflow for the extraction of neuronal signals. The multi-view projections of 3D calcium imaging data undergo sequential processing modules: denoising (Denoise-module), resolution registration (RR-module), background removal (BR-module), and neuronal extraction (NE- module). Subsequently, spatial distances and temporal correlations across multi-view projections are utilized for instance grouping, guided by a physics-based optical model, to facilitate accurate 3D localization of neurons.

## Results

### DeepWonder3D workflow for 3D neuronal extraction from multi-view projections

DeepWonder3D is a deep learning-based framework specifically designed to enable rapid and robust 3D neuronal extraction from multi-view projections of 3D calcium imaging data. This workflow consists of five sequential modules: denoising, resolution registration, background removal, neuronal extraction, and multi-view fusion^29^ (Fig. 1d), each addressing unique challenges in processing multi-view neuronal recordings (Supplementary Fig. 1). Together, these modules constitute a robust approach to extract meaningful neuronal signals. The denoising module^30^ of DeepWonder3D is based on DeepCAD^31,32^, a real-time image enhancement method specifically designed for neuronal imaging. The denoising module removes imaging noise, enhancing the robustness of downstream localization and extraction^33^. To ensure the applicability of the entire analysis workflow across diverse 3D calcium imaging systems^34^, we introduced a Resolution Registration module (RR-module, Supplementary Fig. 2). The RR-module utilizes a hybrid architecture, consisting of convolutional layers, linear upsampling layers, and a U-Net framework, specifically designed to register the resolution of low-resolution images. To train the RR-module, we downsampled original neuron recordings to simulate degraded inputs and optimize the reconstruction based on high-resolution ground truth (Supplementary Fig. 1). This step leverages complementary information across multiple views to achieve both consistent spatial details and inter-view coherence, which are factors essential for resolving complex neuronal structures (Supplementary Fig. 3).

After resolution registration, we employed a Background Removal module (BR-module) for eliminating background signals resulting from scattered photons in 1p imaging, consequently yielding uncontaminated temporal traces of calcium signals. The BR-module consists of a generator and discriminator built upon a 3D U-Net architecture, thereby making full use of the temporal information within neuroimaging videos. The BR-module maps the input data which includes neuronal and background signals onto a video output that preserves only the neuronal calcium signals (Supplementary Fig. 1). The Neuronal extraction module (NE-module) extracts individual neurons from the recording after background signal removal. This module utilizes a 2D U-Net architecture^35^, processes temporally overlapping neuron video chunks, and generates neuronal extraction information (Supplementary Fig. 1). After obtaining initial extraction masks, the NE-module uses spatiotemporal connectivity analysis to connect neuron segments across frames, treating different extractions with substantial spatial overlap as a single neuron. Finally, we used the Multi-View Fusion module (MVF-module) to integrate neuronal spatial localization information from multi-view projections to estimate their 3D localization information and calcium signal temporal activity traces. We performed clustering based on the Pearson correlation of their neuronal temporal traces and the spatial proximity of their localization information. We calculated the final neuron’s 3D position based on a physical optics model of the PSF, while simultaneously fusing the temporal traces within each cluster (Supplementary Fig. 2). By fusing spatiotemporal localization information from multi-view projections, the MVF-module achieves 3D reconstruction of neuronal dynamics, minimizing errors caused by inter-view discrepancies (Supplementary Fig. 4).

Training the DeepWonder3D framework necessitates extensive labeled data. To overcome the challenges and high costs associated with acquiring large-scale manual annotations, we generated simulated data with known ground truth using NAOMi-LF, a tool we developed by extending the Neural Anatomy and Optical Microscopy (NAOMi)^36^ package. Spatially, NAOMi-LF constructs realistic virtual cortical tissue volumes by populating them with fluorescent and non-fluorescent neurons, axons, dendrites, and blood vessels, defining the static ground truth structure (Supplementary Fig. 5a). Temporally, synthetic spike trains generated by a neuronal network model are convolved with models of calcium kinetics and indicator dynamics to produce fluorescence signals for individual neurons (Supplementary Fig. 5b). NAOMi-LF then simulates the complete image formation process: it combines the spatial structure and temporal activity into an instantaneous 3D fluorescence volume, convolves this volume with a view-specific PSF, and incorporates effects like background light scattering (via an occlusion mask) and additive imaging noise to produce a final simulated image. Iteration of this process over time yields the complete simulated multi-view recordings (Supplementary Fig. 5c). This simulation-based training strategy enhances the generalizability of the DeepWonder3D framework across diverse imaging conditions.

### Performance evaluation of DeepWonder3D through simulated light field neuronal recordings

We utilized the NAOMi-LF simulator to generate the results of 1p light field neuroimaging (Fig. 2a), while concurrently obtaining precise neuron 3D spatial localization information and calcium signal temporal activity information. We simulated 20 sets of imaging results to conduct a comprehensive validation of DeepWonder3D; each dataset has a pixel size of 1 μm, a volume size of 500×500×100 μm, and a temporal length of 2000 frames. The final results showed that DeepWonder3D efficiently removed background noise (Fig. 2b), precisely localized neurons in 3D space, and accurately extracted their temporal traces (Fig. 2c, Supplementary Fig. 6). In comparison to MesoLF^37^, DeepWonder3D demonstrated better performance in neuronal extraction, manifested by a superior F1 score (Fig. 2c, Supplementary Fig. 6) and reduced 3D localization errors over all axial shifts^38^ (Fig. 2d). Regarding temporal trace extraction, DeepWonder3D illustrated statistically higher temporal correlation relative to MesoLF (Fig. 2e).

**Figure 2.**
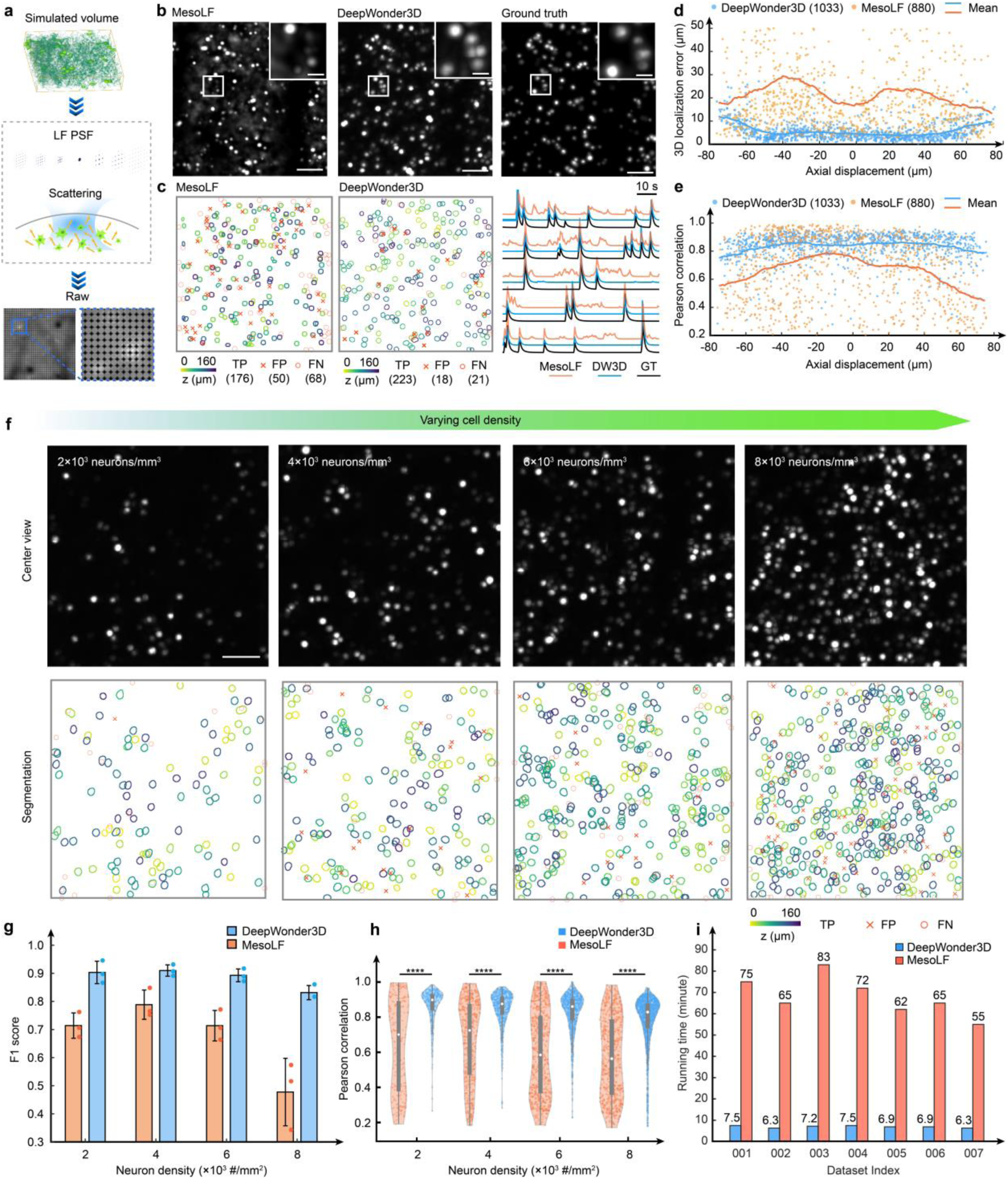
Benchmark of DeepWonder3D with state-of-art algorithms using simulated calcium imaging data. a, Illustration of the simulation data generation process utilizing the NAOMi-LF simulator. Simulated volumes containing neurons with diverse morphological features, including somas, dendrites, axons, and non-fluorescent cells, were subjected to processing through a light field PSF and a scattering model to generate realistic simulated light field images. A magnified view of the simulated light field image is presented in the blue inset box. **b,** Comparative assessment of imaging results processed by MesoLF and DeepWonder3D. The left panel displays the MIP of the standard deviation over time derived from MesoLF reconstruction. The middle panel shows the center-view image after background removal performed by DeepWonder3D. The right panel presents the MIP of the ground truth simulated volume. **c,** Comparison of neuronal extraction outcomes between MesoLF and DeepWonder3D. The left panel illustrates the extraction results obtained using MesoLF. True positive (TP) extractions are indicated by color-coded neuron contours representing their depth within a 160 μm axial range. False positives (FP) are marked with red crosses, and false negatives (FN) are indicated by red circles. The middle panel shows the extraction results from DeepWonder3D with the same labeling convention. The right panel displays example temporal traces of neuronal activity extracted by MesoLF (orange), DeepWonder3D (blue), and the corresponding ground truth traces (black). **d,** Analysis of the 3D localization error for extracted neurons. The 3D localization error is plotted as a function of the neuronal axial displacement for results obtained from MesoLF (orange) and DeepWonder3D (blue). Each data point represents an individual neuron, with the solid lines indicating the mean error at corresponding axial positions. **e,** Pearson correlation analysis of extracted neuronal traces. The Pearson correlation coefficient of the neuronal traces extracted by MesoLF (orange) and DeepWonder3D (blue) relative to the ground truth is presented. The labeling of data points corresponds to that in panel **d**. **f,** Results demonstrating the performance of DeepWonder3D under varying neuronal densities. The top row displays the center-view images after background removal by DeepWonder3D at increasing neuron densities. The bottom row shows the corresponding extraction results from DeepWonder3D for each density, using the same labeling convention as in panel **c**. **g,** Comparison of F1 scores for MesoLF (orange) and DeepWonder3D (blue) across different neuronal densities. F1 scores quantify the accuracy of neuronal extraction. Mean F1 scores and standard deviations are reported for each method at densities of 2, 4, 6, 8×10^3^ neurons mm^-3^ are 0.71±0.05, 0.79±0.05, 0.71±0.05, and 0.48±0.12, respectively. F1 scores of DeepWonder3D with neuron density of 2, 4, 6, 8 ×10^3^ neurons mm^- 3^ are 0.90±0.04, 0.91±0.02, 0.89±0.02, and 0.83±0.03, respectively. Statistical scores are shown in mean ±s.d. across n = 3 simulated recordings for each neuron density. Height of bars: mean. Error bars: s.d. Colored dots: individual measurements (n = 3 simulated recordings). **h,** Pearson correlation of extracted neuronal traces with ground truth under distinct neuronal densities for MesoLF (orange) and DeepWonder3D (blue). Mean Pearson correlation coefficients and standard deviations are provided for each method at densities of 2, 4, 6, 8 ×10^3^ neurons mm^-3^ are 0.63 ±0.22, 0.70 ±0.15, 0.60 ±0.25, and 0.62 ±0.20, respectively. F1 scores of DeepWonder3D with neuron density of 2, 4, 6, 8 ×10^3^ neurons mm^-3^ are 0.84 ±0.09, 0.84 ±0.09, 0.82 ±0.11, and 0.80 ±0.12, respectively. ****P < 1 ×10^−10^ for all density, two-sided Wilcoxon signed-rank test. White circles denote the median, thick gray vertical lines represent the interquartile range, and thin vertical lines indicate upper and lower proximal values. Transparent colored dots show individual data points, and transparent violin-shaped areas illustrate the kernel density estimate of the data distribution. **i**, Comparison of processing running time between MesoLF (orange) and DeepWonder3D (blue). Scale bars: 100 μm (**b**, **f**), 20 μm (zoom-in panel of **b**).

We generated datasets with densities spanning 2, 4, 6, and 8 ×10^3^ neurons mm^-3^, simulating three independent recordings (*n* = 3) for each density level (Fig. 2f, Supplementary Fig. 7). Quantitative analysis focused on two key performance metrics: neuronal extraction accuracy and temporal fidelity of extracted activity traces, comparing DeepWonder3D against the MesoLF method. For neuronal extraction, quantified by the F1 score, DeepWonder3D consistently outperformed MesoLF across all tested densities (Fig. 2g). Specifically, DeepWonder3D achieved mean F1 scores of 0.90±0.04, 0.91±0.02, 0.89±0.02, and 0.83± 0.03 (mean ± standard deviation, s.d.) for densities of 2, 4, 6, and 8 × 10^3^ neurons mm^-3^, respectively. In contrast, MesoLF yielded lower F1 scores of 0.71±0.05, 0.79±0.05, 0.71± 0.05, and 0.48±0.12 under the same conditions. Similarly, when evaluating the temporal accuracy of extracted calcium traces via Pearson correlation with the ground truth signals, DeepWonder3D demonstrated significantly superior performance (Fig. 2h). The temporal correlations achieved by DeepWonder3D were 0.84 ± 0.09, 0.84 ± 0.09, 0.82 ± 0.11, and 0.80 ± 0.12 (mean ± s.d.) for the respective densities. These values were markedly and significantly higher than those obtained with MesoLF (0.63 ± 0.22, 0.70 ± 0.15, 0.60 ± 0.25, and 0.62 ±0.20, mean ±s.d.; ****P < 1 ×10^−10^ for all densities, two-sided Wilcoxon signed-rank test). Furthermore, DeepWonder3D achieves a 10-fold speedup over MesoLF, with speed improvements ranging from +88.5% to +91.3% across seven datasets (Fig. 2i). These results demonstrated that DeepWonder3D maintained high performance and robustness in both accurate neuron identification and faithful extraction of temporal dynamics across a range of neuronal densities, surpassing the comparative baseline method under these challenging simulated conditions.

### Assessing DeepWonder3D performance with a hybrid two-photon/light-field imaging system

Beyond simulated datasets, rigorous validation on real-world experimental data is essential to establish the practical utility of DeepWonder3D. To this end, we performed comprehensive evaluations using *in vivo* recordings acquired from a custom-built hybrid imaging system integrating light field (LF) and 2p microscopy^39^ (Fig. 3a; Methods). This setup allowed us to leverage the high spatial resolution and established signal fidelity of 2p calcium imaging as a functional ground truth for validating the performance of our LF analysis workflow^40^. Meticulous lateral and axial alignment between the two modalities was ensured through fluorescent bead calibration and affine transformation mapping, guaranteeing accurate spatiotemporal correspondence between the LF field-of-view (FOV) and the 2p FOV during simultaneous recordings (Supplementary Note 1).

**Figure 3.**
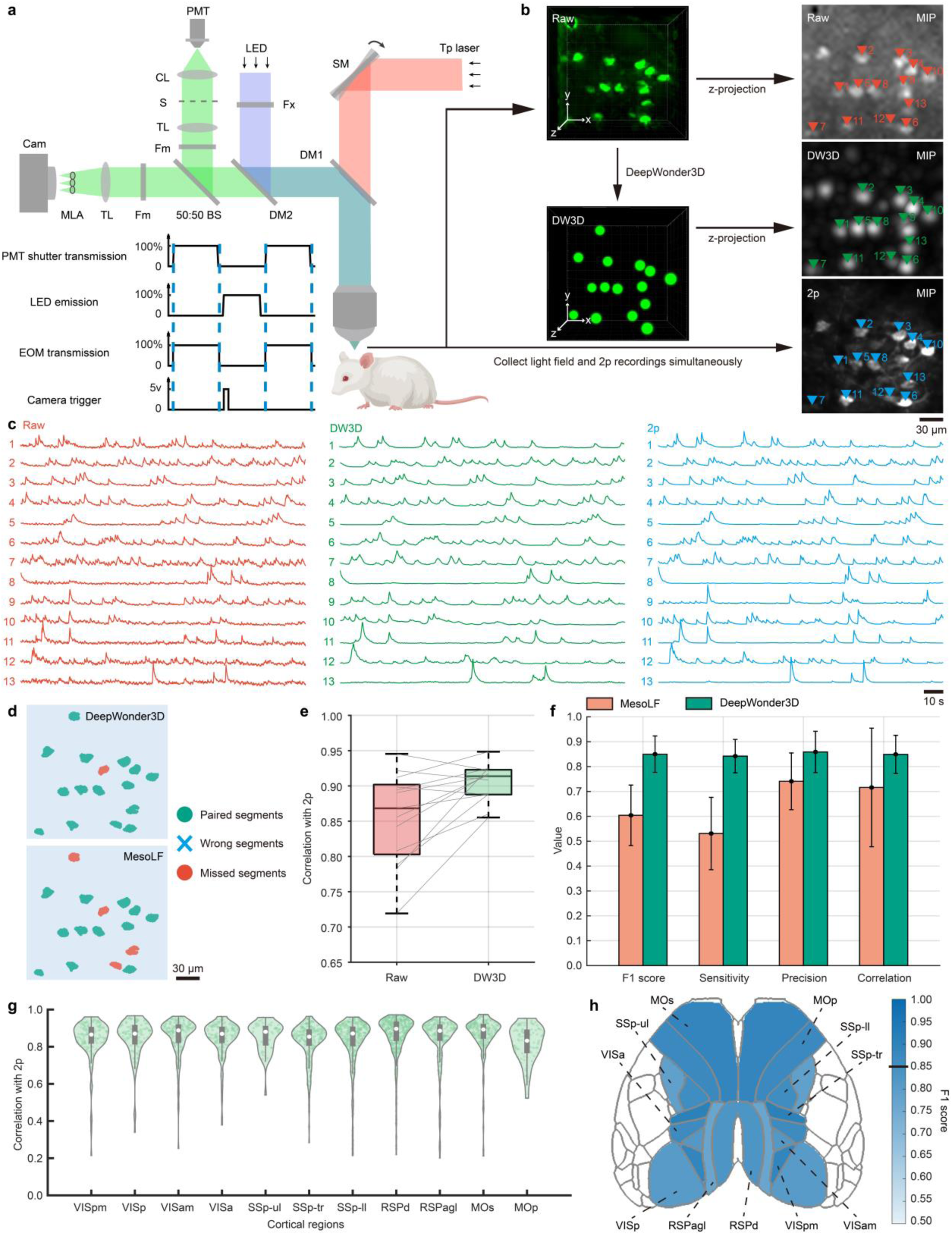
Experimental evaluation of DeepWonder3D using a hybrid two-photon/light- field imaging system. a, Schematic diagram of the custom-built hybrid 2p/LF microscope system. LED, light-emitting diode light source; Tp laser, titanium:sapphire laser; M, mirror; DM, dichroic mirror; BS, beam splitter; Fm, emission filter; Fx, excitation filter; CL, collection lens; TL, tube lens; S, triggerable shutter; CAM, sCMOS camera; MLA, micro-lens array; PMT, photomultiplier tube; SM, galvo scanning mirror; Obj, objective. The inset box on the right illustrates the control signals for coordinating the shutter, LED, electro-optic modulator (EOM), and camera exposure during simultaneous LF and 2p data acquisition. **b,** MIPs of representative neuronal activity recordings. The top panel shows the MIP of the raw light field recording. The middle panel presents the MIP of the light field recording processed by DeepWonder3D. The bottom panel displays the MIP of the simultaneously acquired two- photon recording, which serves as the ground truth. Neurons are indicated by colored triangles. **c,** Extracted calcium traces from the neurons indicated in panel **b**, comparing the three imaging modalities: raw light field (red), DeepWonder3D-processed light field (green), and two-photon (blue). These traces demonstrate the temporal fidelity of the signals obtained after processing with DeepWonder3D. **d,** Comparison of neuronal extraction results from DeepWonder3D (top) and the baseline MesoLF method (bottom) applied to the data shown in panel **b**. Green indicates correctly matched segments corresponding to ground truth neurons, red indicates false negatives, and blue crosses indicate false positives. **e,** Temporal correlations of the calcium traces for the 13 representative neurons shown in panel **c**. The box plot illustrates the distribution of Pearson correlation coefficients. The Pearson correlations between the raw light field and two-photon data, and between the DeepWonder3D-processed light field and two-photon data, are 0.85 ± 0.07 and 0.90 ± 0.03 (mean ± s.d.), respectively. Statistical significance was assessed using a two-sided Wilcoxon signed-rank test (\*\**P* = 0.009). The central black mark represents the median, the left and right edges of the box indicate the 25th and 75th percentiles, and the whiskers extend to the extreme points excluding outliers (defined as 1.5 times the interquartile range). **f,** Quantitative comparison of performance metrics (F1 score, sensitivity, precision, and temporal correlation) for extraction results obtained from MesoLF(orange) and DeepWonder3D (green) across 40 paired 2p/LF recordings. Error bars represent the standard deviation across trials. DeepWonder3D achieved mean F1 score, sensitivity, precision, and temporal correlation values of 0.85 ±0.07, 0.84 ±0.07, 0.86 ±0.08, and 0.85 ±0.08 (mean ±s.d.), respectively. MesoLF yielded corresponding values of 0.60 ± 0.12, 0.53 ± 0.15, 0.74 ± 0.11, and 0.72 ± 0.24 (mean ± s.d.). Statistical significance was determined using a two-sided Wilcoxon signed-rank test: F1 score (****P = 3.57 × 10⁻⁸), sensitivity (****P = 3.57 × 10⁻⁸), precision (****P = 7.05 × 10⁻⁷), temporal correlation (****P < 1 ×10^−10^). **g,** Distribution of temporal correlation coefficients between DeepWonder3D-extracted neuronal traces and the corresponding 2p ground truth across 11 distinct mouse brain regions. The analyzed regions include: posteromedial visual area (VISpm), primary visual area (VISp), anteromedial visual area (VISam), anterior area (VISa), dorsal part of the retrosplenial area (RSPd), lateral agranular part of the retrosplenial area (RSPagl), primary somatosensory areas for the upper limb (SSp-ul), trunk (SSp-tr), and lower limb (SSp- ll), as well as the secondary motor area (MOs) and primary motor area (MOp). **h,** Spatial distribution of neuronal extraction accuracy, represented by F1 scores, achieved by DeepWonder3D. The F1 scores are overlaid on the Allen Common Coordinate Framework (CCF) atlas. The black line on the colorbar indicates the mean F1 score. Scale bars: 30 μm (**b**, **d**), 10 s (**c**).

Qualitative assessment revealed that DeepWonder3D effectively processed the LF recordings, yielding neuronal spatial footprints and temporal calcium dynamics that closely mirrored those obtained from the concurrent high-resolution 2p imaging^41^ (Fig. 3b, c). Visual comparison further suggested superior spatial localization accuracy compared to the baseline MesoLF method (Fig. 3d), and the extracted temporal traces exhibited markedly higher correlation with the 2p ground truth than the unprocessed raw LF signals (Fig. 3e). To quantitatively substantiate these observations, we conducted a detailed performance comparison between DeepWonder3D and MesoLF using a cohort of 40 paired 2p/LF recordings (Fig. 3f). DeepWonder3D demonstrated a statistically significant and substantial advantage in overall neuronal extraction accuracy, achieving a mean F1 score of 0.85 ± 0.07 (mean ±s.d., *n* = 40), compared to 0.60 ± 0.12 obtained with MesoLF (****P = 3.57 × 10⁻⁸, two-sided Wilcoxon signed-rank test). Critically, this enhanced extraction accuracy did not arise from a trade-off favoring either sensitivity or precision, but rather resulted from balanced and significant improvements in both metrics. DeepWonder3D yielded a sensitivity of 0.84 ± 0.07 compared to MesoLF’s 0.53 ± 0.15 (****P = 3.57 × 10⁻⁸) and a precision of 0.86 ± 0.08 compared to MesoLF’s 0.74 ± 0.11 (****P = 7.05 × 10⁻⁷), indicating its superior capability in identifying true positive neurons while concurrently minimizing false positive extractions. In terms of temporal fidelity, DeepWonder3D achieved a significantly higher temporal correlation with the 2p ground truth (0.85 ± 0.08) than MesoLF (0.72 ± 0.24, ****P < 1 ×10^−10^), effectively preserving the underlying calcium dynamics. Furthermore, we assessed the spatial robustness of DeepWonder3D across diverse cortical areas. Analysis across 11 distinct mouse brain regions revealed that DeepWonder3D consistently extracted neuronal activity with high temporal correlation relative to the 2p ground truth (Fig. 3g) and maintained high extraction accuracy (F1 score) across these varied anatomical contexts (Fig. 3h). This underscored the broad applicability and reliability of DeepWonder3D for analyzing LFM data acquired from different cortical areas.

### Depth-dependent performance validation via 3D two-photon imaging

A crucial validation step involves assessing the performance of DeepWonder3D along the axial dimension, given LFM’s volumetric acquisition capability and the inherent focal-plane limitation of standard two-photon microscopy. To establish a depth-resolved ground truth, we integrated an electrically tunable lens (ETL) into the two-photon arm of our hybrid imaging system^42^. This modification enabled rapid axial scanning, allowing us to acquire high-resolution 2p functional data at multiple distinct focal planes (specifically at 50 μm, 75 μm, 100 μm, and 125 μm depth) within the volume simultaneously imaged by the LF modality.

We then compared the neuronal signals extracted by DeepWonder3D from the volumetric LF data against the corresponding depth-specific 2p ground truth recordings. Our results demonstrated that DeepWonder3D accurately captured neuronal identities and their temporal activities across the different focal planes we investigated (Fig. 4a). Specifically, when we compared performance against the MesoLF method, we found that DeepWonder3D achieved higher F1 scores (Fig. 4b) at all tested depths. In our analysis of temporal fidelity, we found that DeepWonder3D maintained a high mean temporal correlation (exceeding 0.87) with the 2p ground truth across all depths, outperforming MesoLF in this metric as well (Fig. 4c). These depth-resolved validation experiments on LF data, utilizing ETL-assisted 2p imaging for ground truthing, further substantiated the robust performance and axial accuracy of our DeepWonder3D workflow. Furthermore, we benchmarked DeepWonder3D against Suite2p^43^ and CNMF-E^44^ on ETL 2p data. While the F1 scores of Suite2p and CNMF-E consistently declined from ∼0.6 as imaging depth increased, DeepWonder3D maintained high and robust performance (F1 scores > 0.8) across all depths (Fig. 4d). In terms of processing speed, DeepWonder3D processed each 1200-frame 3D data (all depths) in just 62.8 seconds on average, substantially faster than Suite2p (225.7 s) and CNMF-E (270.0 s) (Fig. 4e). These findings highlight the rapid and robust capabilities of DeepWonder3D, not only on light-field datasets but also on ETL-based 2p volumes, underscoring its versatility across diverse 3D imaging modalities.

**Figure 4.**
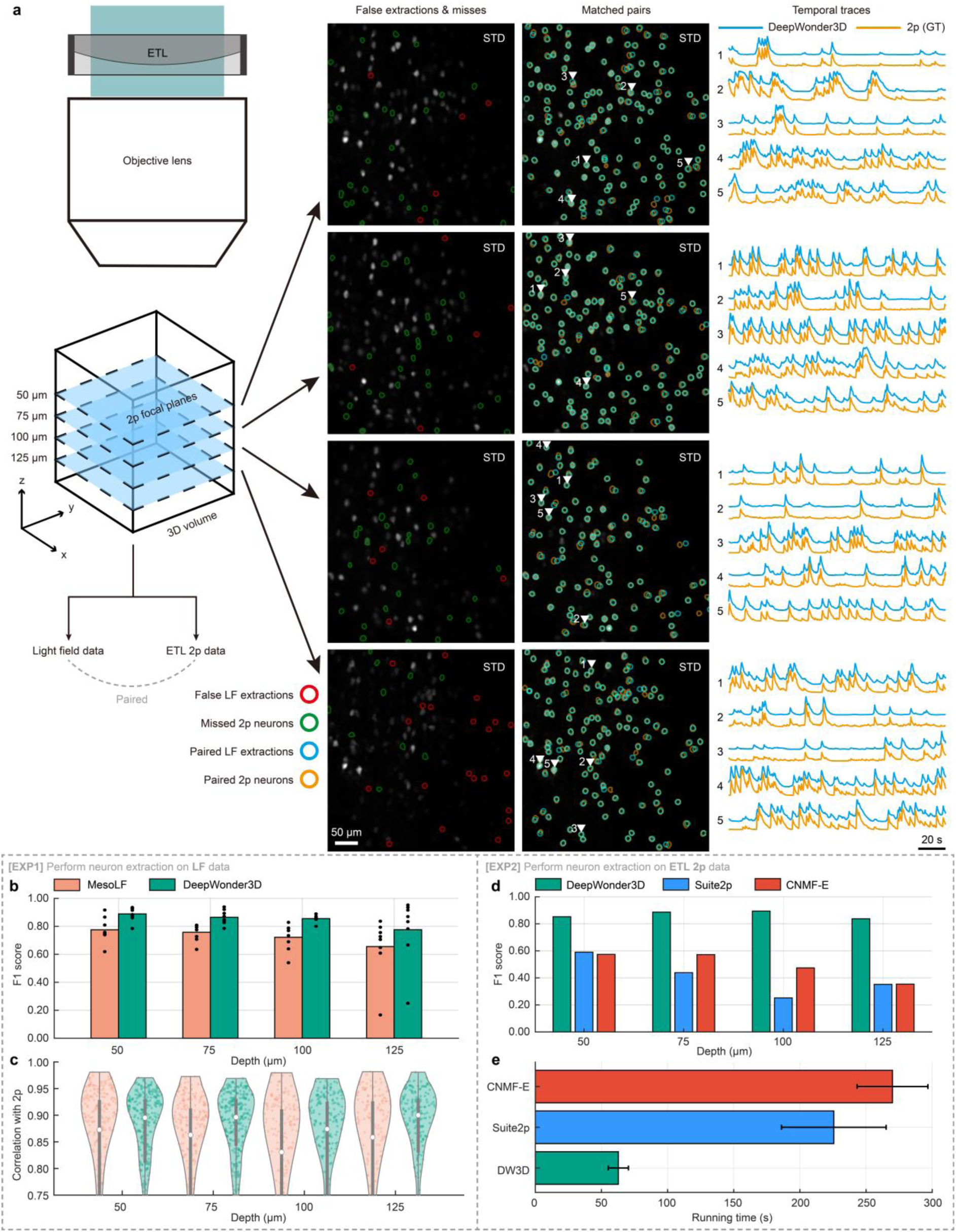
Evaluation of 3D performance using an electrically tunable lens during 2p imaging. a, Representative fields of view acquired simultaneously with LFM and axially scanned two-photon microscopy at four focal planes (50, 75, 100 and 125 µm). 2p focal planes are adjusted using an electrically tunable lens (ETL). Each column shows, from left to right: extraction errors relative to the 2p reference (false positives in red, false negatives in green), correctly matched neuron pairs (DeepWonder3D in cyan, 2p reference in orange), and ΔF/F traces of the matched neurons (same color code). The images are overlaid on the temporal- standard-deviation (temporal-STD) projection of the corresponding 2p recording. **b-c**, Quantitative comparison of neuron extraction on LF data between DeepWonder3D (green) and MesoLF (orange). **b**, F1 score. Bars denote the mean. **c**, Pearson correlation between light field- derived and 2p traces (one point per neuron; *n* = 215, 212, 154 and 134 neurons for the four depths). Violin plots show the kernel density estimate; the white circles indicate the median; the thick gray lines show the interquartile range. **d-e**, Quantitative comparison of neuron extraction on ETL 2p data between DeepWonder3D (green), Suite2p (blue), and CNMF-E (red). **d**, F1 score. Bars denote the mean. **e,** Mean processing time per 1200-frame 3D data (all depths together). Scale bars: 50 µm (images) and 20 s (traces).

To promote the further development of spatiotemporal 3D analysis of neurons, we collected and compiled the light field neuronal imaging data and the registered two-photon neuronal imaging data into the LF2Pneuron dataset. The LF2Pneuron dataset contains 280.72 GB of data, and we have annotated the spatial information of neurons based on the two-photon imaging results.

#### DeepWonder3D as a user-oriented toolbox for large-scale optical neuronal imaging

Recent advancements in microscopy are pushing the frontiers of neuroimaging, enabling visualization of neuronal activity across unprecedented scales and depths. For example, 2pSAM^12^ greatly enhances imaging depth and resolution within scattering brain tissue by computationally integrating multi-view projections, and RUSH3D mesoscope^13^ enables high- throughput imaging with high spatiotemporal resolution across large volumes (e.g., 8,000×6,000×400 µm³at 20 Hz). These two systems enable 3D volume reconstruction by combining multi-view projections through a deconvolution algorithm integrated with Digital Adaptive Optics^45^ (DAO, Supplementary Fig. 4). However, the terabyte-scale, high-resolution datasets generated by these systems present considerable computational challenges, demanding automated, efficient, and robust analysis workflows to fully realize their scientific potential^46^. Existing processing tools often face limitations related to speed, usability, and adaptability to diverse data types and quality levels.

To address this critical need, we refined and validated the multimodal applicability of DeepWonder3D on the 2pSAM system and its high-throughput applicability on the RUSH3D system. Optimized using TensorRT for rapid neural network inference, DeepWonder3D provides an end-to-end solution, transforming raw imaging data into extracted neuronal signals through automated steps including denoising, neuronal extraction, and signal extraction (Fig. 5a). A key strength lies in its versatility; DeepWonder3D demonstrates robust performance across various optical modalities, including conventional 1p, 2p, widefield, and LFM, while effectively handling datasets with both high and low signal-to-noise ratios (SNR). Crucially, its architecture, as demonstrated below, is well-suited for processing challenging data from advanced techniques like 2pSAM and RUSH3D mesoscope.

**Figure 5.**
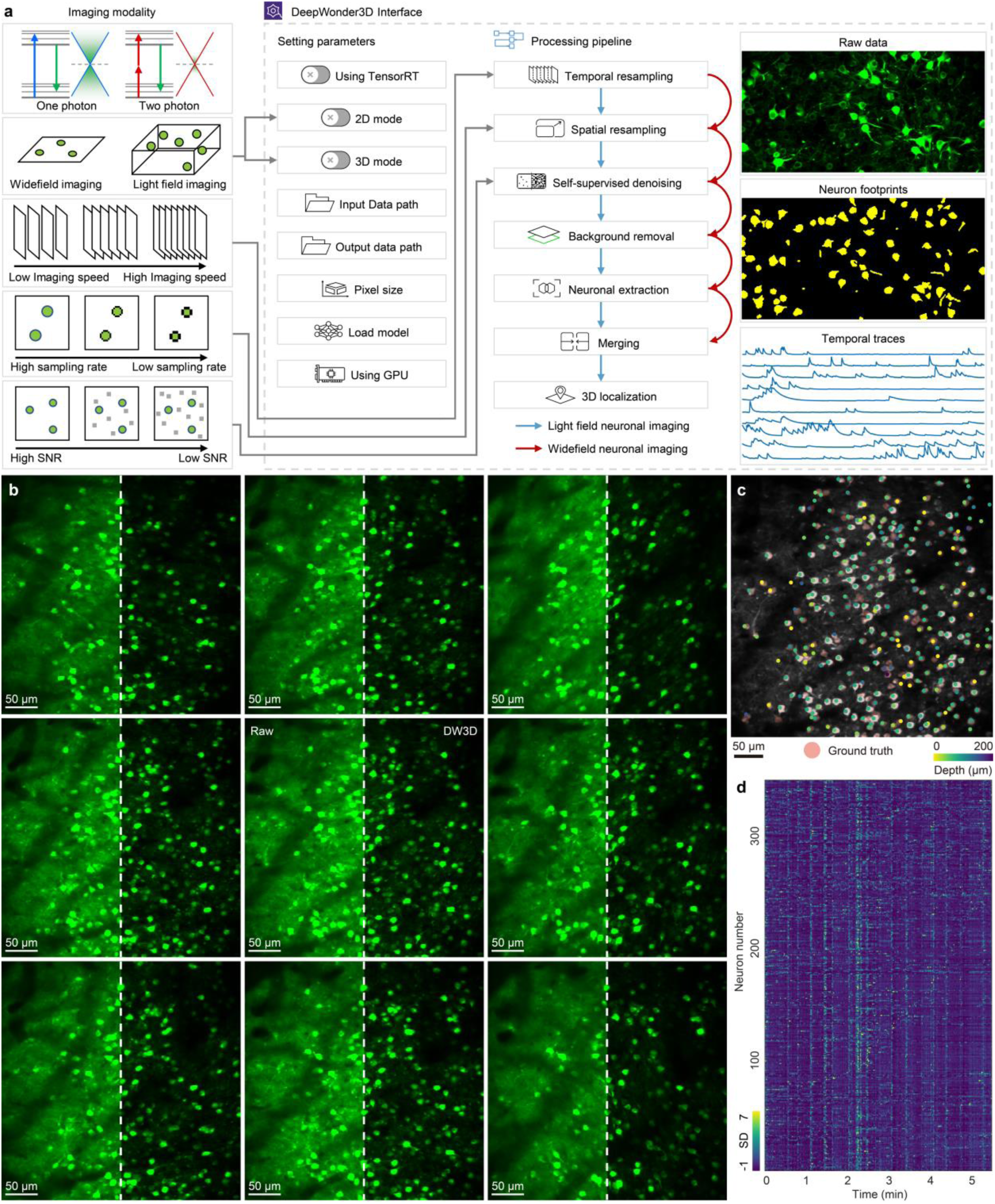
Experimental validation of DeepWonder3D on two-photon synthetic aperture microscopy. **a**, Schematic of the DeepWonder3D processing workflow. Users configure sequential modules (self-supervised denoising, spatiotemporal resampling, background removal, neuronal extraction and 3D localization) and enable GPU or TensorRT acceleration to process both wide-field and light field data across varying sampling rates and signal-to-noise ratios. **b**, Validation on multi-view two-photon calcium imaging data: temporal-STD projections from multiple perspectives (surrounding the central view) are shown before (left panels) and after (right panels) DeepWonder3D processing. **c**. 3D reconstruction of neuron positions derived from the data in **b**. Depth-encoded centroids of all extracted neurons (colored points) are overlaid on the temporal-STD projection of central view; red outlines indicate manually annotated ground-truth neuron locations. **d**. Heatmap of ΔF/F traces for every neuron extracted by DeepWonder3D over the 5-minute recording. Each row represents the activity of a single neuron. Scale bars: 50 μm (**b, c**).

We first validated DeepWonder3D using data acquired via 2pSAM, which captured deep- tissue neuronal activity from 13 distinct viewpoints across a 500 µm×500 µm field of view (FOV) with 1 µm lateral resolution (Fig. 5b). Leveraging its integrated image enhancement and 3D extraction modules, DeepWonder3D accurately identified neuronal locations within the reconstructed volume (Fig. 5c), achieving a high F1 score of 0.88 when compared against expert manual annotations. Furthermore, it reliably extracted time-resolved calcium activity traces for the identified neurons (Fig. 5d), showcasing its utility for functional analysis of deep-brain circuits imaged with multi-view techniques (Supplementary Fig. 8).

To further demonstrate the capability of DeepWonder3D for processing large-scale, high-resolution datasets acquired *in vivo*, we employed the RUSH3D system to record neuronal calcium transients across a substantial cortical volume in an awake, head-fixed mouse. The experimental paradigm involved presenting the mouse with full-field drifting gratings oriented in six distinct directions (0°to 150°in 30°increments) to evoke visual responses (Fig. 6a). This experiment generated a terabyte-scale dataset comprising 3,000 light field frames captured over a 5-minute period.

**Figure 6.**
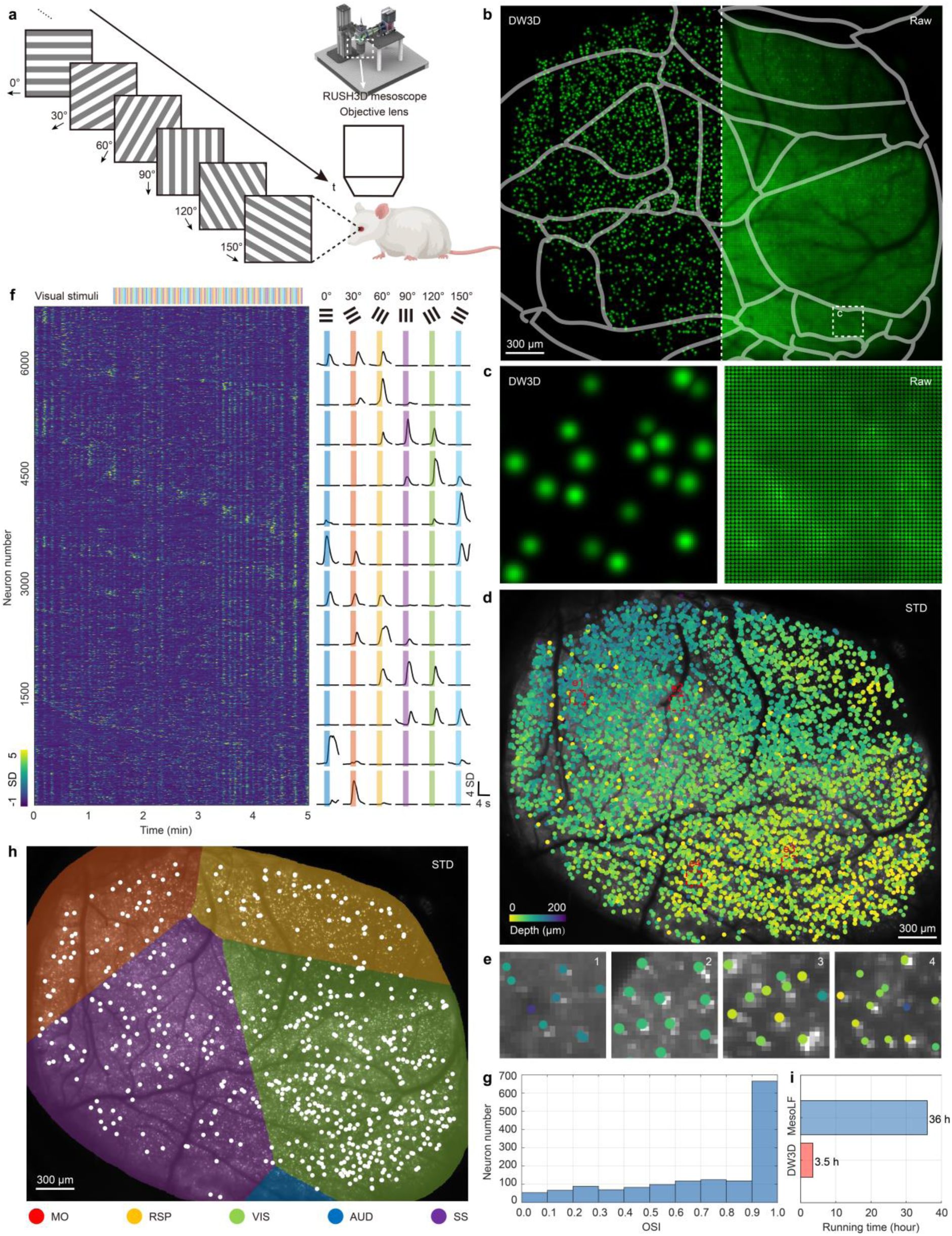
Rapid 3D extraction of large-scale neuronal dynamics in mouse cortex during visual stimulus. a, Experimental configuration. A head-fixed mouse viewed full-field drifting gratings in six directions (0°-150°, 30°steps; arrows indicate drift direction), while cortical activity was recorded via RUSH3D system. **b,** Comparison of raw light field image (left) and DeepWonder3D output of center view (right); cortical area outlines are superimposed in gray. **c,** Zoomed-in region (white dashed box in **b**) of the raw image (left) and corresponding DeepWonder3D output of center view (right). **d**, 3D reconstruction of all extracted neurons (points), color-coded by depth (z-axis), overlaid on the temporal-STD projection of the raw center view. **e**, Detailed views of neurons in the regions marked by red boxes in **d**. **f,** Left, heatmap of ΔF/F traces for more than 6,000 neurons extracted over a 5-minute recording with visual stimuli (colored bars). Right, example ΔF/F traces from representative neurons, with visual stimuli of the six grating directions (colored bars, same color code). **g**, Histogram of OSI values for neurons that showed significant responses to the drifting gratings (two-sided Wilcoxon signed-rank test, *P < 0.05). **h**, Spatial distribution of orientation-selective neurons (OSI > 0.9; *n* = 666), overlaid on the raw temporal-STD projection. Cortical areas are outlined: somatomotor (MO, red), retrosplenial (RSP, yellow), visual (VIS, green), auditory (AUD, blue) and somatosensory (SS, purple). **i**, Computational throughput comparison: mean processing time per 3,000-frame dataset for MesoLF versus DeepWonder3D. Scale bars: 300 μm (**b, d, h**).

We applied the DeepWonder3D workflow to this challenging dataset. First, the preprocessing of DeepWonder3D effectively removed scattering-induced background fluorescence, leading to a marked improvement in the signal-to-noise ratio compared to the raw light field images (Fig. 6b, 6c). After neuronal extraction and multi-view fusion, DeepWonder3D reliably resolved the 3D spatial distribution of active neurons across the cortical depths (Fig. 6d). The high temporal standard deviation (STD) values at the identified neuronal positions confirmed that DeepWonder3D accurately pinpointed locations of prominent neuronal activity in the data (Fig. 6e). Temporally, DeepWonder3D extracted the ΔF/F calcium dynamics for over 6,000 individual neurons from the 5-minute recording, with many traces exhibiting distinct orientation tuning in response to the presented visual stimuli (Fig. 6f). To quantify the orientation tuning properties of the extracted neurons, we calculated the orientation selectivity index (OSI) for all neurons exhibiting significant stimulus-evoked responses^47^ (two-sided Wilcoxon signed-rank test, *P < 0.05), and our analysis revealed that most of them displayed strong orientation tuning. The distribution of OSI values was skewed towards higher selectivity, resulting in a mean OSI of 0.7 across the responsive neurons (Fig. 6g). Notably, we identified 666 neurons exhibiting high orientation selectivity (OSI > 0.9). Mapping the spatial distribution of these highly selective neurons demonstrated a clear concentration within the visual cortical region (Fig. 6h). This localization is consistent with the visual nature of the experimental stimuli, providing physiological validation for the quality of neuronal activity traces extracted by DeepWonder3D. Furthermore, DeepWonder3D demonstrated notable computational efficiency. It processed the entire dataset within 3.5 hours, a processing time 10.3 times faster than that required by MesoLF for the same dataset (Fig. 6i). On an even larger-scale dataset, a 20-minute recording with an 8 mm ×6 mm cortical field of view, DeepWonder3D’s performance was further validated, achieving similarly robust results (Supplementary Fig. 9). Taken together, these results demonstrated the DeepWonder3D workflow’s capacity for rapid and robust analysis of imaging data spanning different scales and potentially diverse modalities.

## Discussion

We introduced a method called DeepWonder3D, a general end-to-end pipeline for rapid and robust 3D neuronal extraction. DeepWonder3D aims to provide a user-oriented and practical workflow to tackle the challenges of imaging noise, scattering, background signals, system diversity, and large-scale data involved in 3D neuronal extraction from multi-view projections of 3D calcium imaging data^48^. Instead of processing voxel by voxel, DeepWonder3D directly leverages the rich information embedded in the multi-view projections. DeepWonder3D sequentially applies self-supervised denoising, imaging resolution registration, simulation- supervised background signal removal, and spatiotemporal information fusion for neuronal extraction. Finally, DeepWonder3D performs multi-view fusion based on optical physical information, leading to the 3D spatial localization and temporal signal extraction of neurons. DeepWonder3D effectively mitigates the impact of noise and scattering on 3D neuronal localization within the workflow, demonstrating better performance compared to advanced methods like MesoLF. DeepWonder3D maintains high neuronal extraction accuracy (F1 score) across varying depths and neuronal densities, while also retaining high temporal fidelity (Pearson correlation). DeepWonder3D achieves a tenfold increase in processing speed compared to MesoLF, enabling the analysis of TB-scale datasets from advanced high-throughput neuronal imaging systems such as RUSH3D mesoscope within a practical timeframe. Furthermore, DeepWonder3D has been verified for compatibility with both single-photon (i.e. LFM, RUSH3D mesoscope) and two-photon (i.e. 2pSAM) modalities and is equipped with a user-oriented toolbox and TensorRT acceleration, lowering the barriers for neurophysiology laboratories to utilize large-scale volumetric imaging capabilities^49^. Consequently, DeepWonder3D provides a general solution for rapid robust high-fidelity 3D neuronal extraction, paving the way for more complex studies of distributed neural circuit activity and behavioral relationships^50^.

While DeepWonder3D has demonstrated many advantages, it also has some potential limitations. Firstly, the BR module of DeepWonder3D depends on training data generated by the NAOMi-LF simulator. Although NAOMi-LF creates training data with various resolutions and signal-to-noise ratios, discrepancies inevitably exist compared to actual collected calcium imaging data. DeepWonder3D’s generalization capability ensures its robust performance when applied to real-world data with various imaging modalities; however, further pushing the precision of digital twins of brain tissue would enhance the progress of simulation-guided methods^51^. With the enhancement of GPU computational capabilities, DeepWonder3D will incorporate transformer-based modules^52^, utilizing attention mechanisms to improve the precision of neuron analysis^53^. Moreover, during the final fusion step, DeepWonder3D simplifies the PSF to a straight line for back-projection, accelerating computation, but it remains to be investigated whether a more precise position estimation method exists. For example, using lightweight neural networks to combine neuronal position information from different angles with the physical information of imaging to derive the final 3D coordinates.

In the future, DeepWonder3D has considerable potential for advancement in improving the landscape of computational science in support of neuroscience^54^. First, the integration of DeepWonder3D with cloud computing could enable real-time 3D localization and analysis of neurons^55^, increasing the efficiency of behavioral experiments in neuroscience^56,57^. By refining the network architecture and fusion modules, the analytical scope could be broadened beyond neuronal cell bodies to investigate activities within finer subcellular structures such as dendrites and axons. The multifunctionality of DeepWonder3D further suggests that it has the potential to be utilized across various model organisms for 3D neuronal imaging, including zebrafish larvae and Caenorhabditis elegans, as well as 3D organoid models^58^. Additionally, as voltage imaging becomes widely adopted in neuroscience, DeepWonder3D is anticipated to extend its capabilities to analyze neural voltage imaging data^59^, thereby comprehensively covering all imaging modalities, labeling techniques, and model organisms in neuronal imaging^60^.

## Methods

### Hybrid 2p/LF ground truth recordings

To validate our algorithms in achieving correct neuronal activities, we built a joint 2p and LF detection system. The schematic of the custom-built 2p microscope is shown in Supplementary Fig. 10. A titanium-sapphire laser system (MaiTai HP, Spectra-Physics) served as the 2p excitation source (920 nm central wavelength, pulse width <100 fs, 80 MHz repetition rate). A half-wave plate (AQWP10M-980, Thorlabs) and an EOM (350-80LA-02, Conoptics) modulated the excitation power. The scanned beam went through a scan lens (SL50-2P2, Thorlabs) and a tube lens (TTL200MP, Thorlabs) and an objective (20×/0.45 NA, UCPLFLN20X, Olympus). A high-precision piezo actuator (P-725, Physik Instrumente) drove the objective for fast axial scanning. To extend the depth of field of the 2p excitation, we reduced the beam size at the back aperture of the objective with an iris. The effective excitation NA was about 0.25 in our imaging experiments, yielding ∼27 µm axial range. A long-pass dichroic mirror (DMLP650L, Thorlabs) was used to separate fluorescence signals from femtosecond laser beam by reflecting the fluorescence signals and transmitting the infrared laser light.

For the 1p LF excitation path, a long-pass dichroic (DMLP505L, Thorlabs) in the original detection path of TPLSM was used to send 470 nm-centered LED light (M470L4-C1 and MF475-35, Thorlabs) to the objective. To jointly record LF excitation and 2p excitation, a 50:50 (reflectance:transmission) nonpolarizing plate beam splitter (BSW27, Thorlabs) was placed after the second dichroic to separate fluorescent signals for PMT (PMT1001, Thorlabs) and camera (Panda 4.2, Pco) with micro lens array (F number is 20, pitch size is 97.5 μm), respectively. A pair of fluorescence filters (MF525-39, Thorlabs; ET510/80M, Chroma) was configured in front of both the PMT and the camera to fully block both femtosecond laser and

LED excitation beam. The back aperture of the objective was optically conjugated to the detection surface of the PMT with a 4f system (TTL200-A and AC254-050-A, Thorlabs).

To avoid excitation crosstalk and protect PMT from high-flux widefield emission photons, we added a linear galvo that served as an optical shutter for the PMT detection path, which deflected LF fluorescent photons when LED was on. We further configured the EOM to be blocked during LF imaging. The LED (M470L4-C1) was in trigger mode with a typical rising and falling time less than 1 ms, with further reduced duration time to avoid PMT overexposure (Supplementary Fig. 10).

### Structure of DeepWonder3D

#### Denoising module

The denoising module adopts a 3D U-Net^61^ architecture to effectively enhance the signal- to-noise ratio of input recordings. Trained with L1 and L2 reconstruction losses, it minimizes the pixel-wise difference between the denoised output and ground truth, preserving fine details while suppressing noise to ensure structural and temporal integrity of neuronal signals.

#### Resolution registration module

The resolution registration module is a composite architecture designed to upscale and refine images through the integration of multiple computational layers. This module comprises three convolutional layers, a linear upscaling layer, and a U-Net architecture, arranged sequentially to capture both local and global spatial features for high-resolution image reconstruction. It takes as input single-frame images from multiple views, leveraging their complementary information to enhance spatial details and ensure inter-view consistency after resolution registration. During training, the raw recordings serve as the ground truth (GT), while their downsampled counterparts are used as input, simulating the degradation process.

The model is trained to optimize a combination of L1 and L2 losses, promoting both pixel- wise accuracy and overall structural similarity between the predicted and ground truth images.

#### Background removal module

This module aims to remove background signals from input recordings, isolating neuronal firing activity. The simulation process can obtain not only a normal video 𝑎, which includes both background signals and neuronal activity, but also a corresponding background-free video 𝑏, containing only neuron firing. In DeepWonder3D, the background removal module employs a Generative Adversarial Network (GAN) composed of two 3D U-Net^61^ architectures: a generator 𝐺 and a discriminator 𝐷. During training, the generator learns to map 𝑎 to 𝑏, guided by a loss function that combines L1 and L2 losses: 𝓛_𝓖_ = 𝑳_𝟏_(𝒃, 𝑮(𝒂)) + 𝑳_𝟐_(𝒃, 𝑮(𝒂)) . Meanwhile, the discriminator is trained to distinguish between real background-free videos and those generated by 𝐺, using a classical GAN loss 𝓛_𝓓_ = 𝐥 𝐧(𝑫(𝒃)) + 𝐥 𝐧 (𝟏 − 𝑫(𝑮(𝒂))). In the inference phase, only the generator 𝐺 is utilized to process input recordings, producing background-free outputs that retain only the neuronal signals.

#### Neuronal extraction module

The neuronal extraction module is built upon a 3D U-Net^61^ architecture. A simulated background-free recording is split into 32-frame patches with a 16-frame overlap between consecutive patches. The patches are processed with CaImAn^62^ to generate 2D neuron masks. During training, the 32-frame patches are used as inputs, while the corresponding 2D neuron masks are treated as the ground truth. The loss function for the neuronal extraction network is a combination of L1 and L2 losses. After processing each patch of the video, spatiotemporal connectivity analysis is performed to link the mask sequences and merge overlapping segments. Neurons that overlap spatially but are temporally separated (for instance, when neuron segments appear in different frames) are treated as distinct candidates. This process enables the extraction of individual neurons by compiling a list of candidates from the entire recording.

#### Multi-view fusion module

The multi-view fusion module integrates results from multi-view projections to compute the final 3D coordinates and temporal traces for each neuron. This process involves several key steps: First, the Pearson correlation between the temporal traces of neurons across different views is computed, and candidates with high correlation are clustered together. Next, within each cluster, spatial clustering is performed based on Euclidean distance. If multiple sub- clusters are extracted, only the largest one is retained to avoid errors from incorrect candidates. Then, clusters with fewer than four candidates are discarded. Finally, the 3D coordinates of each neuron are estimated based on a physics-based optical model given the PSF, while the temporal trace of each neuron is obtained by averaging the temporal traces of the candidates within each cluster.

### Simulation

#### Realistic data simulation

To create realistic cortical tissues and simulate multi-view projections of 3D calcium imaging data, we employed NAOMi-LF, a tool we developed by extending the Neural Anatomy and Optical Microscopy (NAOMi)^36^ package. Spatially, NAOMi-LF constructs realistic virtual cortical tissue volumes by populating them with fluorescent and non- fluorescent neurons, axons, dendrites, and blood vessels, defining the static ground truth structure (Supplementary Fig. 5a). Temporally, synthetic spike trains generated by a neuronal network model are convolved with models of calcium kinetics and indicator dynamics to produce fluorescence signals for individual neurons (Supplementary Fig. 5b). NAOMi-LF then simulates the complete image formation process: it combines the spatial structure and temporal activity into an instantaneous 3D fluorescence volume, convolves this volume with a view- specific PSF, and incorporates effects like background light scattering (via an occlusion mask) and additive imaging noise to produce a final simulated image. Iteration of this process over time yields the complete simulated multi-view recordings (Supplementary Fig. 5c).

#### Noise simulation

Imaging sensors (e.g., sCMOS, CMOS, and CCDs) exhibit variations in quantum efficiency (QE) and noise response. Noise is also influenced by the expression levels of calcium indicators in neurons. The number of fluorescence photons produced in a unit area of the sample can be expressed as^63^:

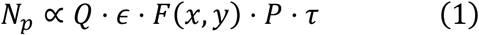

Here, 𝑄 represents the quantum efficiency of fluorophores with an extinction coefficient 𝜖, 𝐹(𝑥, 𝑦) is the fluorophore concentration at a given position, 𝑃 denotes the power density, and 𝜏 is the integration time of the camera. The sensor signal can be further described as^64^:

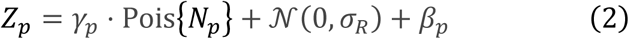

where 𝛾_𝑝_ is a gain factor applied to the Poisson distribution (Pois), 𝛽_𝑝_is the offset introduced during analog-to-digital (AD) conversion, and 𝒩(0, 𝜎_𝑅_) represents Gaussian-distributed readout noise with a mean of zero and a standard deviation 𝜎_𝑅_. For a typical sCMOS sensor, the parameters are approximately γ_𝑝_ ≈ 2.2, β_𝑝_ ≈ 100, and σ_𝑅_ ≈ 200 (ref. ^65^).

### Performance metrics

#### Correlation score

Pearson’s correlation coefficient is used as the temporal metric to assess the similarity between inferred neuronal activities and ground truths. The ground truth activities were available for simulation data, while for joint 2p/LF validation data, the ground truth activities were obtained by running CaImAn^62^ on 2p datasets (Supplementary Note 1).

#### Neuronal extraction scores

To evaluate neuronal extraction performance, it is essential to acquire ground truth extractions. For simulated data, the ground truths were predetermined and readily available. For real data, the ground truths were obtained from matched and aligned 2p data, where neurons were manually annotated based on their spatial locations and activity patterns. Specifically, neuron candidates were selected from correlation and standard deviation images of the raw 2p recordings, focusing on those that stood out from the background and matched the typical neuron size (10∼15 µm in diameter). Candidates with weak or noisy activity were then excluded after reviewing the raw recording. Finally, each neuron was annotated using ImageJ’s ROI manager, and the zipped ROIs were imported into Python for evaluation.

Once ground truth annotations were obtained, a specially designed Python script was used to automatically evaluate extraction performance based on specific criteria. An extraction is classified as a true positive (TP) if its correlation with the corresponding ground truth neuronal signal exceeds 0.2 and the distance between their centroids is less than 20 μm. Otherwise, the extraction is classified as a false positive (FP). Ground truths that are not identified by the algorithm are considered false negatives (FN). Segmentation accuracy is quantified using the F1-score, defined as the harmonic mean of precision and recall.

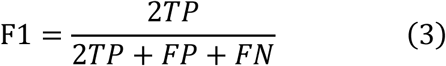

The extraction precision score is defined as

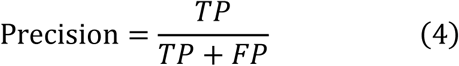

The extraction sensitivity score is defined as

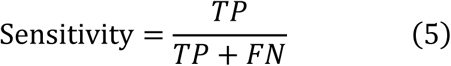

#### Orientation selectivity score

To quantify the degree to which individual neurons exhibit preferential responses to specific stimulus orientations, we calculated the Orientation Selectivity Index (OSI), which is defined as^66^

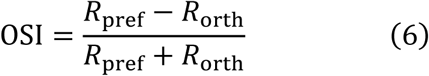

where 𝑅_pref_is the response at the preferred direction and 𝑅_orth_ is the response at the direction orthogonal to the preferred one. A higher OSI value indicates greater selectivity for orientation. Although the OSI could be calculated for all neurons, we restricted our analysis to include only those neurons demonstrating statistically significant responses to the drifting gratings (two- sided Wilcoxon signed-rank test, *P < 0.05). Neurons with non-significant responses were excluded.

## Data availability

We have no restriction on data availability. All source data have been archived and made publicly available at https://doi.org/10.5281/zenodo.15383434. The dataset has been split into multiple parts, with different versions of this archive containing different parts.

## Code availability

Source codes, executable software, and all the other related resources are readily accessible on our GitHub page https://github.com/yujiachenjerry/DeepWonder3D.

## Acknowledgments

This work was supported by grants from the National Key Research and Development Program of China (grant nos. 2023YFC3402600).

## Author contributions

Q.D., R.H. and J.W. conceived and led the project, conceptualized the DeepWonder3D workflow, guided data analysis, and wrote the manuscript. Y.C. accelerated and optimized the DeepWonder3D workflow, performed simulations, compiled the LF2Pneuron dataset (including preprocessing and annotations), analyzed data, and wrote the manuscript. G.Z. designed and implemented the DeepWonder3D workflow, performed simulations, and wrote the manuscript. M.W. collected the raw data of LF2Pneuron dataset and analyzed data. Y.Z., J.X., and Z.Z. provided critical support on system setup, imaging procedures, and biological experiments.

## References

1. Zeng, H. & Sanes, J. R. Neuronal cell-type classification: challenges, opportunities and the path forward. Nat. Rev. Neurosci. 18, 530–546 (2017).

2. Stringer, C., Pachitariu, M., Steinmetz, N., Reddy, C. B., Carandini, M. & Harris, K. D. Spontaneous behaviors drive multidimensional, brainwide activity. Science 364, eaav7893 (2019).

3. Panzeri, S., Moroni, M., Safaai, H. & Harvey, C. D. The structures and functions of correlations in neural population codes. Nat. Rev. Neurosci. 23, 551–567 (2022).

4. Demas, J., Manley, J., Tejera, F., Barber, K., Kim, H., Traub, F. M., Chen, B. & Vaziri, A. High-speed, cortex-wide volumetric recording of neuroactivity at cellular resolution using light beads microscopy. Nat. Methods 18, 1103–1111 (2021).

5. Manley, J., Lu, S., Barber, K., Demas, J., Kim, H., Meyer, D., Traub, F. M. & Vaziri, A. Simultaneous, cortex-wide dynamics of up to 1 million neurons reveal unbounded scaling of dimensionality with neuron number. Neuron 112, 1694–1709.e5 (2024).

6. Kim, T. H. & Schnitzer, M. J. Fluorescence imaging of large-scale neural ensemble dynamics. Cell 185, 9–41 (2022).

7. Zhang, Y., Rózsa, M., Liang, Y., Bushey, D., Wei, Z., Zheng, J., Reep, D., Broussard, G. J., Tsang, A., Tsegaye, G., Narayan, S., Obara, C. J., Lim, J.-X., Patel, R., Zhang, R., Ahrens, M. B., Turner, G. C., Wang, S. S.-H., Korff, W. L., Schreiter, E. R., Svoboda, K., Hasseman, J. P., Kolb, I. & Looger, L. L. Fast and sensitive GCaMP calcium indicators for imaging neural populations. Nature 615, 884–891 (2023).

8. Wu, J., Lu, Z., Jiang, D., Guo, Y., Qiao, H., Zhang, Y., Zhu, T., Cai, Y., Zhang, X., Zhanghao, K., Xie, H., Yan, T., Zhang, G., Li, X., Jiang, Z., Lin, X., Fang, L., Zhou, B., Xi, P., Fan, J., Yu, L. & Dai, Q. Iterative tomography with digital adaptive optics permits hour-long intravital observation of 3D subcellular dynamics at millisecond scale. Cell 184, 3318–3332.e17 (2021).

9. Feng, X., Ma, Y. & Gao, L. Compact light field photography towards versatile three-dimensional vision. Nat. Commun. 13, 3333 (2022).

10. Levoy, M., Ng, R., Adams, A., Footer, M. & Horowitz, M. Light field microscopy. ACM Trans Graph 25, 924–934 (2006).

11. Zhang, Y., Lu, Z., Wu, J., Lin, X., Jiang, D., Cai, Y., Xie, J., Wang, Y., Zhu, T., Ji, X. & Dai, Q. Computational optical sectioning with an incoherent multiscale scattering model for light-field microscopy. Nat. Commun. 12, 6391 (2021).

12. Zhao, Z., Zhou, Y., Liu, B., He, J., Zhao, J., Cai, Y., Fan, J., Li, X., Wang, Z., Lu, Z., Wu, J., Qi, H. & Dai, Q. Two-photon synthetic aperture microscopy for minimally invasive fast 3D imaging of native subcellular behaviors in deep tissue. Cell 186, 2475–2491.e22 (2023).

13. Zhang, Y., Wang, M., Zhu, Q., Guo, Y., Liu, B., Li, J., Yao, X., Kong, C., Zhang, Y., Huang, Y., Qi, H., Wu, J., Guo, Z. V. & Dai, Q. Long-term mesoscale imaging of 3D intercellular dynamics across a mammalian organ. Cell 187, 1–19 (2024).

14. Pégard, N. C., Liu, H.-Y., Antipa, N., Gerlock, M., Adesnik, H. & Waller, L. Compressive light-field microscopy for 3D neural activity recording. Optica 3, 517–524 (2016).

15. Wagner, N., Beuttenmueller, F., Norlin, N., Gierten, J., Boffi, J. C., Wittbrodt, J., Weigert, M., Hufnagel, L., Prevedel, R. & Kreshuk, A. Deep learning-enhanced light-field imaging with continuous validation. Nat. Methods 18, 557–563 (2021).

16. Wang, Z., Zhu, L., Zhang, H., Li, G., Yi, C., Li, Y., Yang, Y., Ding, Y., Zhen, M., Gao, S., Hsiai, T. K. & Fei, P. Real-time volumetric reconstruction of biological dynamics with light-field microscopy and deep learning. Nat. Methods 18, 551–556 (2021).

17. Truong, T. V., Holland, D. B., Madaan, S., Andreev, A., Keomanee-Dizon, K., Troll, J. V., Koo, D. E. S., McFall-Ngai, M. J. & Fraser, S. E. High-contrast, synchronous volumetric imaging with selective volume illumination microscopy. Commun. Biol. 3, 1–8 (2020).

18. Quicke, P., Howe, C. L., Song, P., Jadan, H. V., Song, C., Knöpfel, T., Neil, M., Dragotti, P. L., Schultz, S. R. & Foust, A. J. Subcellular resolution three-dimensional light-field imaging with genetically encoded voltage indicators. Neurophotonics 7, 035006 (2020).

19. Hua, X., Han, K., Mandracchia, B., Radmand, A., Liu, W., Kim, H., Yuan, Z., Ehrlich, S. M., Li, K., Zheng, C., Son, J., Silva Trenkle, A. D., Kwong, G. A., Zhu, C., Dahlman, J. E. & Jia, S. Light-field flow cytometry for high-resolution, volumetric and multiparametric 3D sin- gle-cell analysis. Nat. Commun. 15, 1975 (2024).

20. Zhang, Z., Bai, L., Cong, L., Yu, P., Zhang, T., Shi, W., Li, F., Du, J. & Wang, K. Imaging volumetric dynamics at high speed in mouse and zebrafish brain with confocal light field mi- croscopy. Nat. Biotechnol. 39, 74–83 (2021).

21. Broxton, M., Grosenick, L., Yang, S., Cohen, N., Andalman, A., Deisseroth, K. & Levoy, M. Wave optics theory and 3-D deconvolution for the light field microscope. Opt. Express 21, 25418–25439 (2013).

22. Streich, L., Boffi, J. C., Wang, L., Alhalaseh, K., Barbieri, M., Rehm, R., Deivasigamani, S., Gross, C. T., Agarwal, A. & Prevedel, R. High-resolution structural and functional deep brain imaging using adaptive optics three-photon microscopy. Nat. Methods 18, 1253–1258 (2021).

23. Cao, R., Li, Y., Wang, W., Zhang, G., Wang, G., Sun, Y., Ren, W., Sun, J., Hou, Y., Xu, X., Hu, J., Lu, Y., Li, C., Wu, J., Li, M., Qu, J. & Xi, P. Dark-based Optical Sectioning assists Background Removal in Fluorescence Microscopy. 2024.03.02.578598 Preprint at 10.1101/2024.03.02.578598 (2024)

24. Kalantari, N. K., Wang, T.-C. & Ramamoorthi, R. Learning-based view synthesis for light field cameras. ACM Trans Graph 35, 193:1–193:10 (2016).

25. Yang, R., Xiao, T., Cheng, Y., Li, A., Qu, J., Liang, R., Bao, S., Wang, X., Wang, J., Suo, J., Luo, Q. & Dai, Q. Sharing massive biomedical data at magnitudes lower bandwidth using implicit neural function. Proc. Natl. Acad. Sci. 121, e2320870121 (2024).

26. Howe, C. L., Zhao, K. L. Y., Verinaz-Jadan, H., Song, P., Barnes, S. J., Dragotti, P. L. & Foust, A. J. Light-field deep learning enables high-throughput, scattering-mitigated calcium imaging. 2025.03.17.643718 Preprint at 10.1101/2025.03.17.643718 (2025)

27. Gui, J., Chen, T., Zhang, J., Cao, Q., Sun, Z., Luo, H. & Tao, D. A Survey on Self-Su- pervised Learning: Algorithms, Applications, and Future Trends. IEEE Trans. Pattern Anal. Mach. Intell. 46, 9052–9071 (2024).

28. Sekh, A. A., Opstad, I. S., Godtliebsen, G., Birgisdottir, Å. B., Ahluwalia, B. S., Agarwal, K. & Prasad, D. K. Physics-based machine learning for subcellular segmentation in living cells. Nat. Mach. Intell. 3, 1071–1080 (2021).

29. Zhao, J., Fu, Z., Yu, T. & Qiao, H. V2V3D: View-to-View Denoised 3D Reconstruction for Light-Field Microscopy. Preprint at 10.48550/arXiv.2504.07853 (2025)

30. Lehtinen, J., Munkberg, J., Hasselgren, J., Laine, S., Karras, T., Aittala, M. & Aila, T. Noise2Noise: Learning Image Restoration without Clean Data. Preprint at 10.48550/arXiv.1803.04189 (2018)

31. Li, X., Zhang, G., Wu, J., Zhang, Y., Zhao, Z., Lin, X., Qiao, H., Xie, H., Wang, H., Fang, L. & Dai, Q. Reinforcing neuron extraction and spike inference in calcium imaging using deep self-supervised denoising. Nat. Methods 18, 1395–1400 (2021).

32. Li, X., Li, Y., Zhou, Y., Wu, J., Zhao, Z., Fan, J., Deng, F., Wu, Z., Xiao, G., He, J., Zhang, Y., Zhang, G., Hu, X., Chen, X., Zhang, Y., Qiao, H., Xie, H., Li, Y., Wang, H., Fang, L. & Dai, Q. Real-time denoising enables high-sensitivity fluorescence time-lapse imaging be- yond the shot-noise limit. Nat. Biotechnol. 41, 282–292 (2023).

33. Lecoq, J., Oliver, M., Siegle, J. H., Orlova, N., Ledochowitsch, P. & Koch, C. Removing independent noise in systems neuroscience data using DeepInterpolation. Nat. Methods 18, 1401–1408 (2021).

34. Lim, B., Son, S., Kim, H., Nah, S. & Lee, K. M. Enhanced Deep Residual Networks for Single Image Super-Resolution. in 2017 IEEE Conf. Comput. Vis. Pattern Recognit. Workshop CVPRW 1132–1140 (2017). doi:10.1109/CVPRW.2017.151

35. Ronneberger, O., Fischer, P. & Brox, T. U-Net: Convolutional Networks for Biomedical Image Segmentation. in Med. Image Comput. Comput.-Assist. Interv. – MICCAI 2015 (eds. Navab, N., Hornegger, J., Wells, W. M. & Frangi, A. F.) 234–241 (Springer International Pub- lishing, 2015). doi:10.1007/978-3-319-24574-4_28

36. Song, A., Gauthier, J. L., Pillow, J. W., Tank, D. W. & Charles, A. S. Neural anatomy and optical microscopy (NAOMi) simulation for evaluating calcium imaging methods. J. Neu- rosci. Methods 358, 109173 (2021).

37. Nöbauer, T., Zhang, Y., Kim, H. & Vaziri, A. Mesoscale volumetric light-field (MesoLF) imaging of neuroactivity across cortical areas at 18 Hz. Nat. Methods 20, 600–609 (2023).

38. Vogelstein, J. T., Packer, A. M., Machado, T. A., Sippy, T., Babadi, B., Yuste, R. & Paninski, L. Fast Nonnegative Deconvolution for Spike Train Inference From Population Cal- cium Imaging. J. Neurophysiol. 104, 3691–3704 (2010).

39. Denk, W., Strickler, J. H. & Webb, W. W. Two-Photon Laser Scanning Fluorescence Microscopy. Science 248, 73–76 (1990).

40. Svoboda, K., Denk, W., Kleinfeld, D. & Tank, D. W. In vivo dendritic calcium dynamics in neocortical pyramidal neurons. Nature 385, 161–165 (1997).

41. Chen, T.-W., Wardill, T. J., Sun, Y., Pulver, S. R., Renninger, S. L., Baohan, A., Schreiter, E. R., Kerr, R. A., Orger, M. B., Jayaraman, V., Looger, L. L., Svoboda, K. & Kim, D. S. Ultrasensitive fluorescent proteins for imaging neuronal activity. Nature 499, 295–300 (2013).

42. Grewe, B. F., Langer, D., Kasper, H., Kampa, B. M. & Helmchen, F. High-speed in vivo calcium imaging reveals neuronal network activity with near-millisecond precision. Nat. Meth- ods 7, 399–405 (2010).

43. Pachitariu, M., Stringer, C., Dipoppa, M., Schröder, S., Rossi, L. F., Dalgleish, H., Ca- randini, M. & Harris, K. D. Suite2p: beyond 10,000 neurons with standard two-photon micros- copy. 061507 Preprint at 10.1101/061507 (2017)

44. Zhou, P., Resendez, S. L., Rodriguez-Romaguera, J., Jimenez, J. C., Neufeld, S. Q., Gio- vannucci, A., Friedrich, J., Pnevmatikakis, E. A., Stuber, G. D., Hen, R., Kheirbek, M. A., Sabatini, B. L., Kass, R. E. & Paninski, L. Efficient and accurate extraction of in vivo calcium signals from microendoscopic video data. eLife 7, e28728 (2018).

45. Zhao, J., Zhao, Z., Wu, J., Yu, T. & Qiao, H. PNR: Physics-informed Neural Represen- tation for high-resolution LFM reconstruction. Preprint at 10.48550/arXiv.2409.18223 (2024)

46. Abe, T., Kinsella, I., Saxena, S., Buchanan, E. K., Couto, J., Briggs, J., Kitt, S. L., Glass- man, R., Zhou, J., Paninski, L. & Cunningham, J. P. Neuroscience Cloud Analysis As a Service: An open-source platform for scalable, reproducible data analysis. Neuron 110, 2771–2789.e7 (2022).

47. Goris, R. L. T., Simoncelli, E. P. & Movshon, J. A. Origin and function of tuning diversity in macaque visual cortex. Neuron 88, 819–831 (2015).

48. Pnevmatikakis, E. A., Soudry, D., Gao, Y., Machado, T. A., Merel, J., Pfau, D., Reardon, T., Mu, Y., Lacefield, C., Yang, W., Ahrens, M., Bruno, R., Jessell, T. M., Peterka, D. S., Yuste, R. & Paninski, L. Simultaneous Denoising, Deconvolution, and Demixing of Calcium Imaging Data. Neuron 89, 285–299 (2016).

49. Zhang, Y., Zhang, G., Han, X., Wu, J., Li, Z., Li, X., Xiao, G., Xie, H., Fang, L. & Dai, Q. Rapid detection of neurons in widefield calcium imaging datasets after training with syn- thetic data. Nat. Methods 20, 747–754 (2023).

50. Wilkinson, M. D., Dumontier, M., Aalbersberg, Ij. J., Appleton, G., Axton, M., Baak, A., Blomberg, N., Boiten, J.-W., da Silva Santos, L. B., Bourne, P. E., Bouwman, J., Brookes, A. J., Clark, T., Crosas, M., Dillo, I., Dumon, O., Edmunds, S., Evelo, C. T., Finkers, R., Gonza- lez-Beltran, A., Gray, A. J. G., Groth, P., Goble, C., Grethe, J. S., Heringa, J., ’t Hoen, P. A. C., Hooft, R., Kuhn, T., Kok, R., Kok, J., Lusher, S. J., Martone, M. E., Mons, A., Packer, A. L., Persson, B., Rocca-Serra, P., Roos, M., van Schaik, R., Sansone, S.-A., Schultes, E., Sengstag, T., Slater, T., Strawn, G., Swertz, M. A., Thompson, M., van der Lei, J., van Mulli- gen, E., Velterop, J., Waagmeester, A., Wittenburg, P., Wolstencroft, K., Zhao, J. & Mons, B. The FAIR Guiding Principles for scientific data management and stewardship. Sci. Data 3, 160018 (2016).

51. Markram, H., Meier, K., Lippert, T., Grillner, S., Frackowiak, R., Dehaene, S., Knoll, A., Sompolinsky, H., Verstreken, K., DeFelipe, J., Grant, S., Changeux, J.-P. & Saria, A. Intro- ducing the human brain project. Procedia Comput. Sci. 7, 39–42 (2011).

52. Dosovitskiy, A., Beyer, L., Kolesnikov, A., Weissenborn, D., Zhai, X., Unterthiner, T., Dehghani, M., Minderer, M., Heigold, G., Gelly, S., Uszkoreit, J. & Houlsby, N. An Image is Worth 16x16 Words: Transformers for Image Recognition at Scale. in (2020). at https://open-review.net/forum?id=YicbFdNTTy

53. Vaswani, A., Shazeer, N., Parmar, N., Uszkoreit, J., Jones, L., Gomez, A. N., Kaiser, Ł. ukasz & Polosukhin, I. Attention is All you Need. in Adv. Neural Inf. Process. Syst. 30, (Curran Associates, Inc., 2017).

54. Deisseroth, K. Optogenetics: 10 years of microbial opsins in neuroscience. Nat. Neurosci. 18, 1213–1225 (2015).

55. Shang, C.-F., Wang, Y.-F., Zhao, M.-T., Fan, Q.-X., Zhao, S., Qian, Y., Xu, S.-J., Mu, Y., Hao, J. & Du, J.-L. Real-time analysis of large-scale neuronal imaging enables closed-loop investigation of neural dynamics. Nat. Neurosci. 27, 1014–1018 (2024).

56. Kaltenecker, D., Al-Maskari, R., Negwer, M., Hoeher, L., Kofler, F., Zhao, S., Todorov, M., Rong, Z., Paetzold, J. C., Wiestler, B., Piraud, M., Rueckert, D., Geppert, J., Morigny, P., Rohm, M., Menze, B. H., Herzig, S., Berriel Diaz, M. & Ertürk, A. Virtual reality-empowered deep-learning analysis of brain cells. Nat. Methods 21, 1306–1315 (2024).

57. Satyanarayanan, M. The Emergence of Edge Computing. Computer 50, 30–39 (2017).

58. Bassett, D. S. & Sporns, O. Network neuroscience. Nat. Neurosci. 20, 353–364 (2017).

59. Abdelfattah, A. S., Kawashima, T., Singh, A., Novak, O., Liu, H., Shuai, Y., Huang, Y.- C., Campagnola, L., Seeman, S. C., Yu, J., Zheng, J., Grimm, J. B., Patel, R., Friedrich, J., Mensh, B. D., Paninski, L., Macklin, J. J., Murphy, G. J., Podgorski, K., Lin, B.-J., Chen, T.- W., Turner, G. C., Liu, Z., Koyama, M., Svoboda, K., Ahrens, M. B., Lavis, L. D. & Schreiter, E. R. Bright and photostable chemigenetic indicators for extended in vivo voltage imaging. Science 365, 699–704 (2019).

60. Stuart, T., Butler, A., Hoffman, P., Hafemeister, C., Papalexi, E., Mauck, W. M., Hao, Y., Stoeckius, M., Smibert, P. & Satija, R. Comprehensive Integration of Single-Cell Data. Cell 177, 1888–1902.e21 (2019).

61. Çiçek, Ö., Abdulkadir, A., Lienkamp, S. S., Brox, T. & Ronneberger, O. 3D U-Net: Learning Dense Volumetric Segmentation from Sparse Annotation. in Med. Image Comput. Comput.-Assist. Interv. – MICCAI 2016 19th Int. Conf. Athens Greece Oct. 17-21 2016 Proc. Part II 424–432 (Springer-Verlag, 2016). doi:10.1007/978-3-319-46723-8_49

62. Giovannucci, A., Friedrich, J., Gunn, P., Kalfon, J., Brown, B. L., Koay, S. A., Taxidis, J., Najafi, F., Gauthier, J. L., Zhou, P., Khakh, B. S., Tank, D. W., Chklovskii, D. B. & Pnevmatikakis, E. A. CaImAn an open source tool for scalable calcium imaging data analysis. eLife 8, 1–45 (2019).

63. Sandison, D. R. & Webb, W. W. Background rejection and signal-to-noise optimization in confocal and alternative fluorescence microscopes. Appl. Opt. 33, 603–615 (1994).

64. Mandracchia, B., Hua, X., Guo, C., Son, J., Urner, T. & Jia, S. Fast and accurate sCMOS noise correction for fluorescence microscopy. Nat. Commun. 11, 94 (2020).

65. Huang, F., Hartwich, T. M. P., Rivera-Molina, F. E., Lin, Y., Duim, W. C., Long, J. J., Uchil, P. D., Myers, J. R., Baird, M. A., Mothes, W., Davidson, M. W., Toomre, D. & Bew- ersdorf, J. Video-rate nanoscopy using sCMOS camera-specific single-molecule localization algorithms. Nat. Methods 10, 653–658 (2013).

66. Zhao, X., Chen, H., Liu, X. & Cang, J. Orientation-selective Responses in the Mouse Lateral Geniculate Nucleus. J. Neurosci. 33, 12751–12763 (2013).

