## Supplementary Information for "Rapid robust high-fidelity 3D neuronal extraction from multi-view projections"

### Tables of contents

**Supplementary Notes**

| **Supplementary Note 1** | System alignment, motion correction, and ground-truth validation for coupled light field and two-photon imaging |
| --- | --- |

**Supplementary Figures**

| **Supplementary Figure 1** | Training data generation and training process for key modules in DeepWonder3D. |
| --- | --- |
| **Supplementary Figure 2** | Architecture of the resolution registration and 3D localization modules in DeepWonder3D. |
| **Supplementary Figure 3** | Upsampling improves spatial localization accuracy in multi-view calcium imaging. |
| **Supplementary Figure 4** | Principle of aberration correction based on multi-view neuronal consistency. |
| **Supplementary Figure 5** | NAOMi-LF workflow for generating simulated light field and multi-view calcium imaging datasets. |
| **Supplementary Figure 6** | Comparative evaluation of DeepWonder3D and MesoLF on simulated single-photon calcium imaging data. |
| **Supplementary Figure 7** | Performance comparison between DeepWonder3D and MesoLF under varying neuronal densities. |
| **Supplementary Figure 8** | Application of DeepWonder3D to two-photon multi-view *in vivo* calcium imaging data. |
| **Supplementary Figure 9** | Application of DeepWonder3D to large-scale *in vivo* optical neurophysiology data. |
| **Supplementary Figure 10** | Schematic of the hybrid light field and two-photon microscope for simultaneous neuronal activity recording. |

### Supplementary Note 1: System alignment, motion correction, and ground-truth validation for coupled light field and two-photon imaging

**System assembly and focal-plane alignment**

After integrating the light field and two-photon (2p) imaging paths, we coarsely aligned their focal planes using a 20 µm thick planar plant section. The 2p microscope was first focused on the specimen; subsequently, the light field tube lens was adjusted so that the 2p focal plane lay within the light field depth of field. To accommodate the lower native resolution of the light field modality at its focal plane, we maintained a nominal 20 µm offset between the two focal planes during this coarse alignment.

For fine alignment, a mono-layer of 1 µm fluorescent beads were deposited onto a microscope slide. Z-stacks were acquired with both modalities using a piezoelectric objective scanner. The optimal focal depth for each system was determined independently, and the light field tube lens was further adjusted, taking into account the relative system magnifications, to ensure precise co-focusing of both imaging planes.

**Spatial registration via affine transformation**

With the two modalities focused at the same depth, sparse bead images were captured by both systems. Corresponding bead centroids were manually identified across modalities, and an affine transformation mapping the light field field of view (FOV) to the 2p FOV was computed using *imregtform* in MATLAB. This pre-calibrated transformation matrix was applied in subsequent *in vivo* experiments to ensure lateral correspondence between light field and 2p recordings.

**Motion correction**

Each light field and 2p time series was first corrected independently for within‑modality motion artifacts using the NormCorre algorithm¹. The resulting motion‐corrected stacks were then spatially registered to one another by applying the pre-computed affine transformation, thereby producing pairwise‐aligned light field and 2p datasets for downstream analysis.

**Generation of 2p ground truths**

To derive reliable ground-truth neuronal footprints and activity traces, we implemented a semi-automated workflow combining CaImAn² with expert curation. First, the unrestricted neuronal extraction of CaImAn was applied to the 2p volumes to generate candidate spatial masks and associated fluorescence traces. A panel of experienced neuroimaging analysts then reviewed and corrected each mask to produce a validated set of ground‑truth spatial footprints. These curated footprints served as seeds for a second CaImAn run in seeded mode, yielding refined spatial components and their corresponding temporal signals.

**Evaluation of neuronal extraction**

We assessed neuronal extraction performance by pairing each extracted light field neuron with a unique 2p ground-truth neuron. A valid pairing required (1) a centroid separation < 20 µm and (2) a Pearson correlation coefficient between ΔF/F traces > 0.2. If multiple ground-truth neurons met these criteria for a single extracted light field neuron, the highest-correlated 2p neuron was selected. Once paired, both the light field and 2p neurons were removed from further consideration. Unpaired light field extractions were classified as false positives, while unpaired 2p ground-truth neurons were classified as false negatives. Extraction accuracy was quantified by computing precision, sensitivity (recall), and F1 scores. Temporal fidelity was evaluated by calculating the Pearson correlation between the ΔF/F traces of each true positive pair: one inferred from the light field data and the other from the 2p reference (CaImAn).

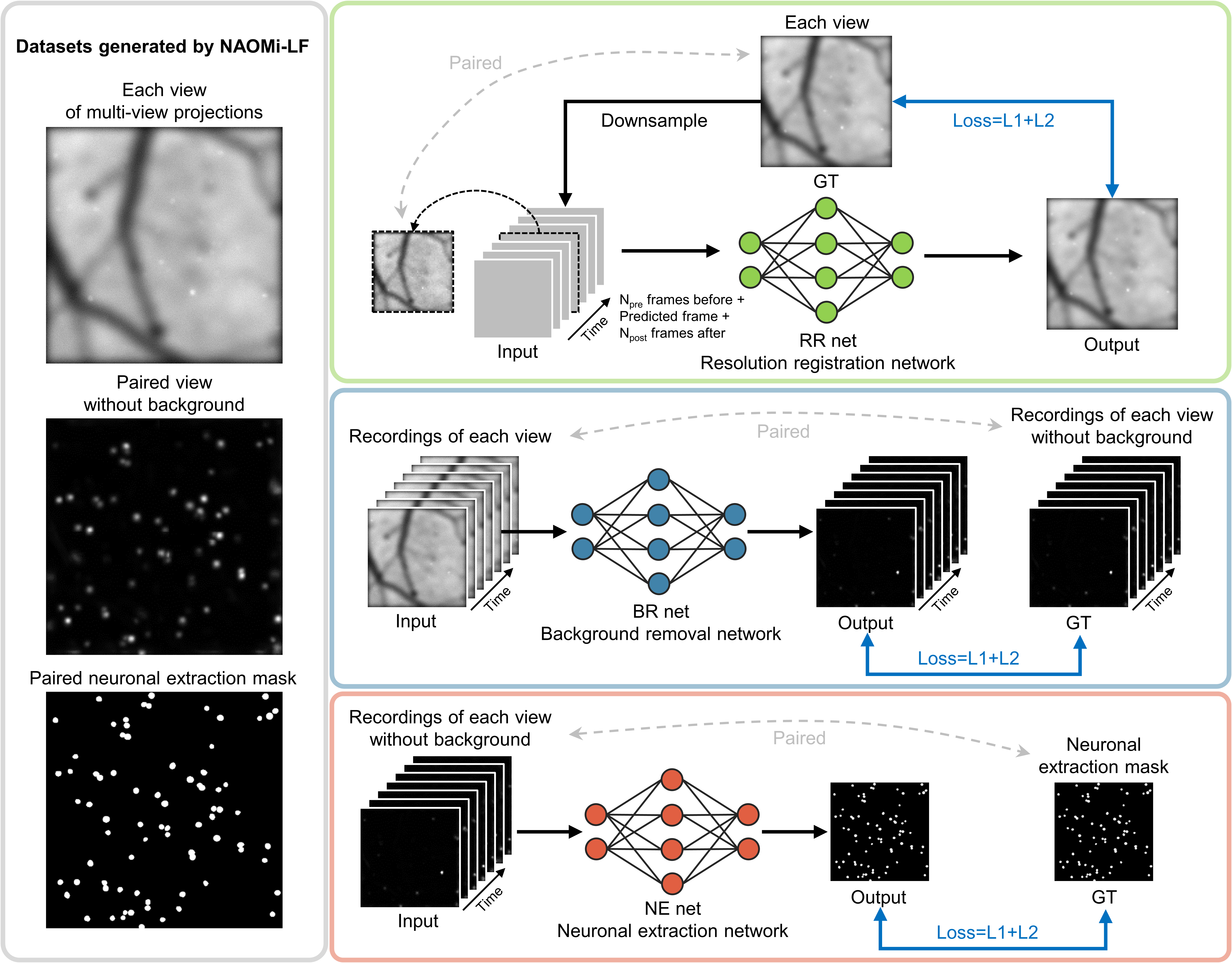

**Supplementary Figure 1 | Training data generation and training process for key modules in DeepWonder3D .** Training data for DeepWonder3D is generated by the NAOMi simulator, which provides synthetic time-series recordings of single-photon calcium imaging, corresponding background-free versions, and ground-truth neuronal spatial localizations. For training the resolution registration (RR) module, the input consists of multi-frame, low-resolution neuronal recordings, while the target is the high-resolution single-frame image at the intermediate time point. This design enabled the model to recover spatial detail through temporal redundancy. The background removal (BR) module is trained using high-resolution recordings as input and corresponding background-free data as the target. Multi-frame input allows the model to leverage spatiotemporal patterns for effective separation of neuronal signals from scattering-induced background. The neuronal extraction (NE) module is trained with high-resolution, background-free recordings as input and neuron spatial annotations as the target. The model learns to identify individual neuronal footprints based on spatial intensity patterns and morphological consistency.

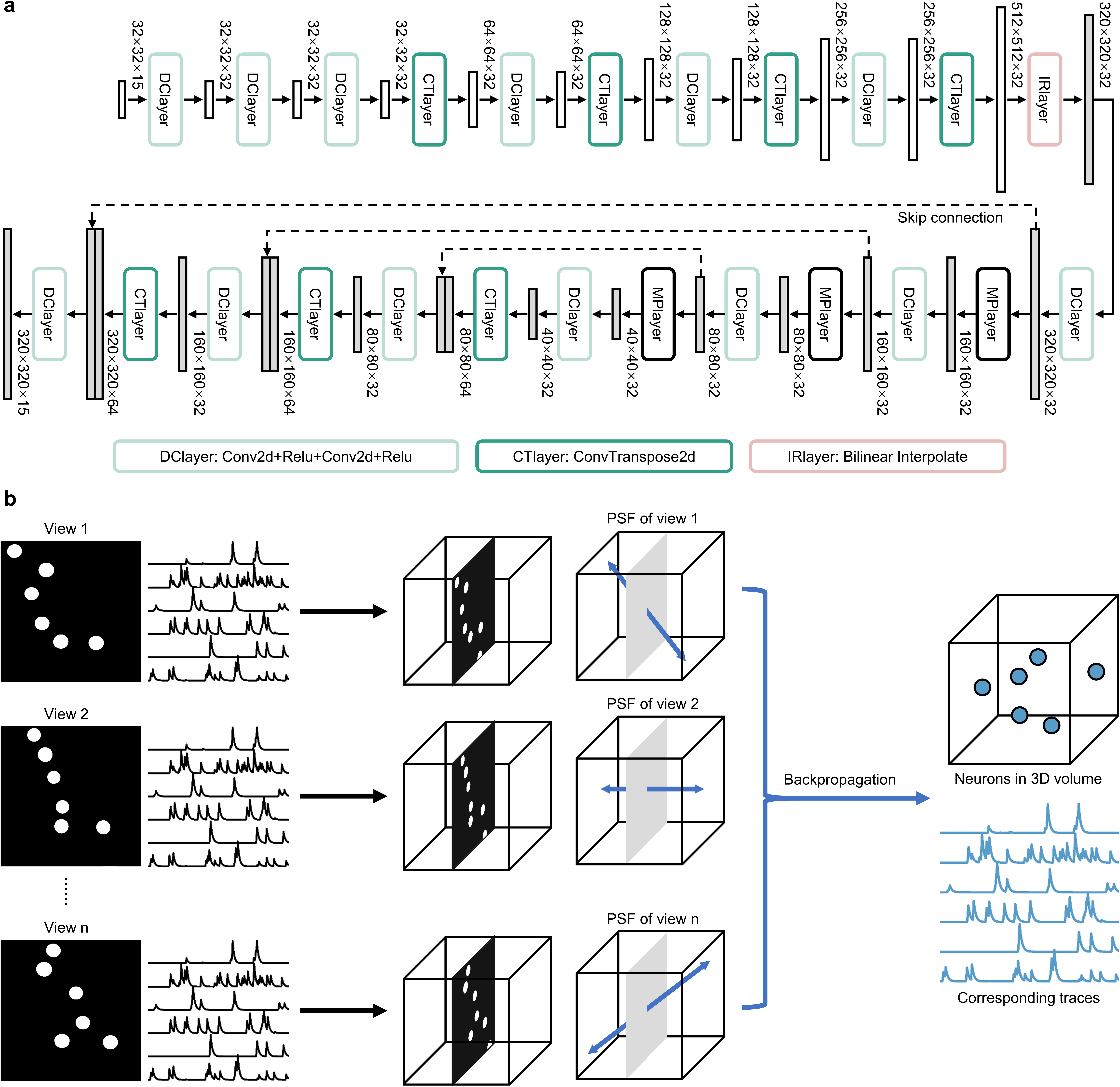
**Supplementary Figure 2 | Architecture of the resolution registration and 3D localization modules in DeepWonder3D . a**, Schematic of the resolution registration module. The input image is first processed by a series of convolutional layers to extract spatial features. These features are then upsampled to restore the original image size. A U-Net architecture is employed at the final stage to enhance fine structural details and spatial coherence. Each sub-module is annotated with its function and name. **b**, Diagram of the 3D neuronal localization module. Neuronal extraction masks are first obtained independently from multiple views. Each view is associated with a specific light-ray propagation direction determined by the PSF of the imaging system. The extracted neuron masks are then back-projected into a shared 3D coordinate space. A voting-based scheme is used to estimate the final 3D location of each neuron by integrating multi-view spatial evidence.

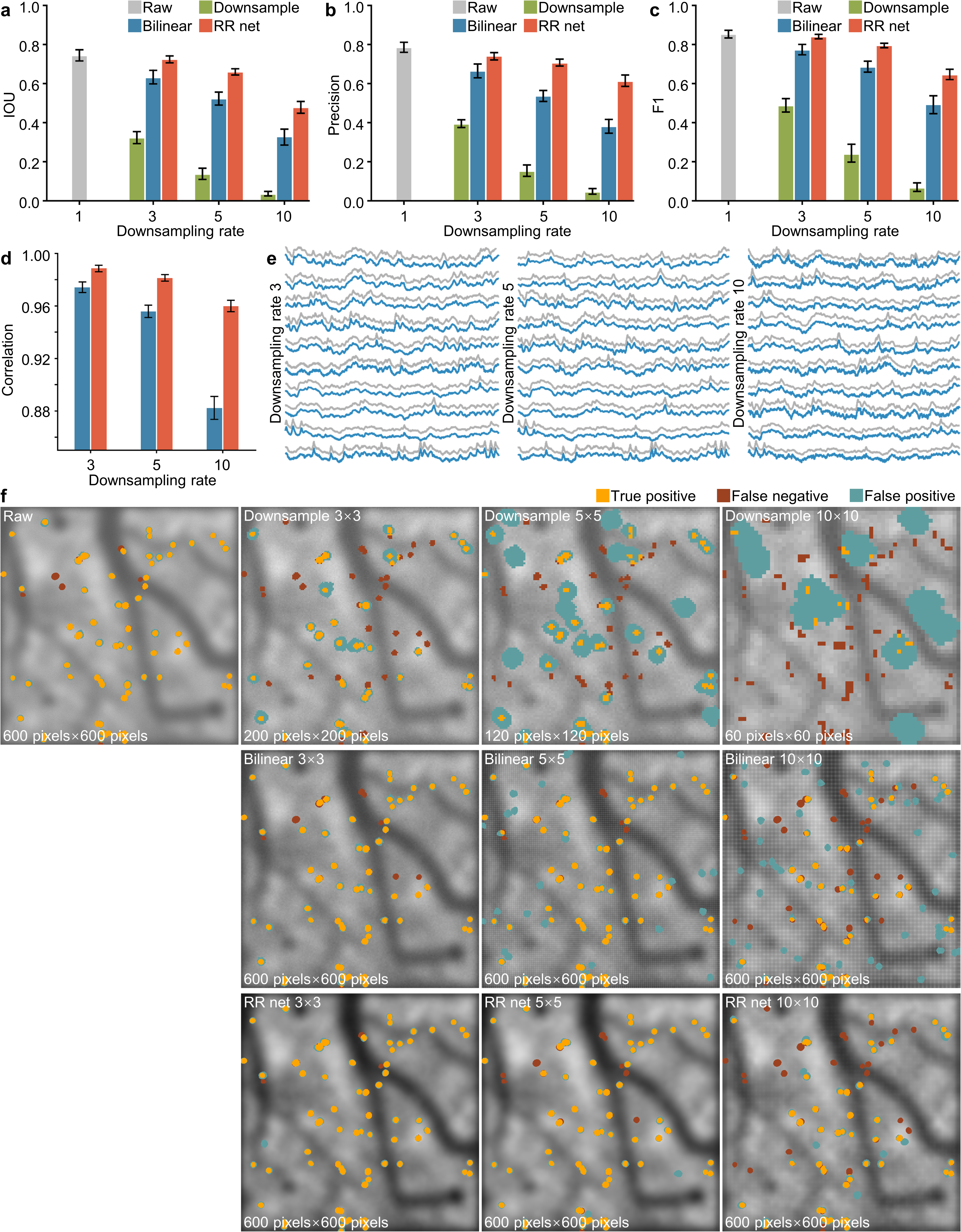

**Supplementary Figure 3 | Upsampling improves spatial localization accuracy in multi-view calcium imaging.** To evaluate the impact of resolution alignment on neuronal localization accuracy, we used simulated single-photon calcium imaging data with 1 μm pixel resolution and corresponding ground-truth spatial annotations. CaImAn was used as the baseline extraction method and applied across four different conditions: original-resolution data, downsampled data, linearly interpolated data, and RR-module enhanced data (produced by the resolution registration module in DeepWonder3D). **a**, Comparison of spatial extraction performance (intersection over union, IoU) across varying downsampling factors and upsampling strategies. **b**, Comparison of extraction precision under the same conditions. **c**, F1 score comparisons indicating overall extraction accuracy. **d**, Temporal fidelity of calcium signal extraction, measured by Pearson correlation with ground-truth fluorescence traces. **e**, Example traces comparing calcium signal reconstruction across different resolutions. **f**, Visualized extraction masks of neuronal footprints under each method. These results demonstrate that learned resolution alignment via RR-module substantially outperforms linear interpolation and mitigates performance degradation caused by spatial downsampling, improving both spatial extraction and temporal signal fidelity.

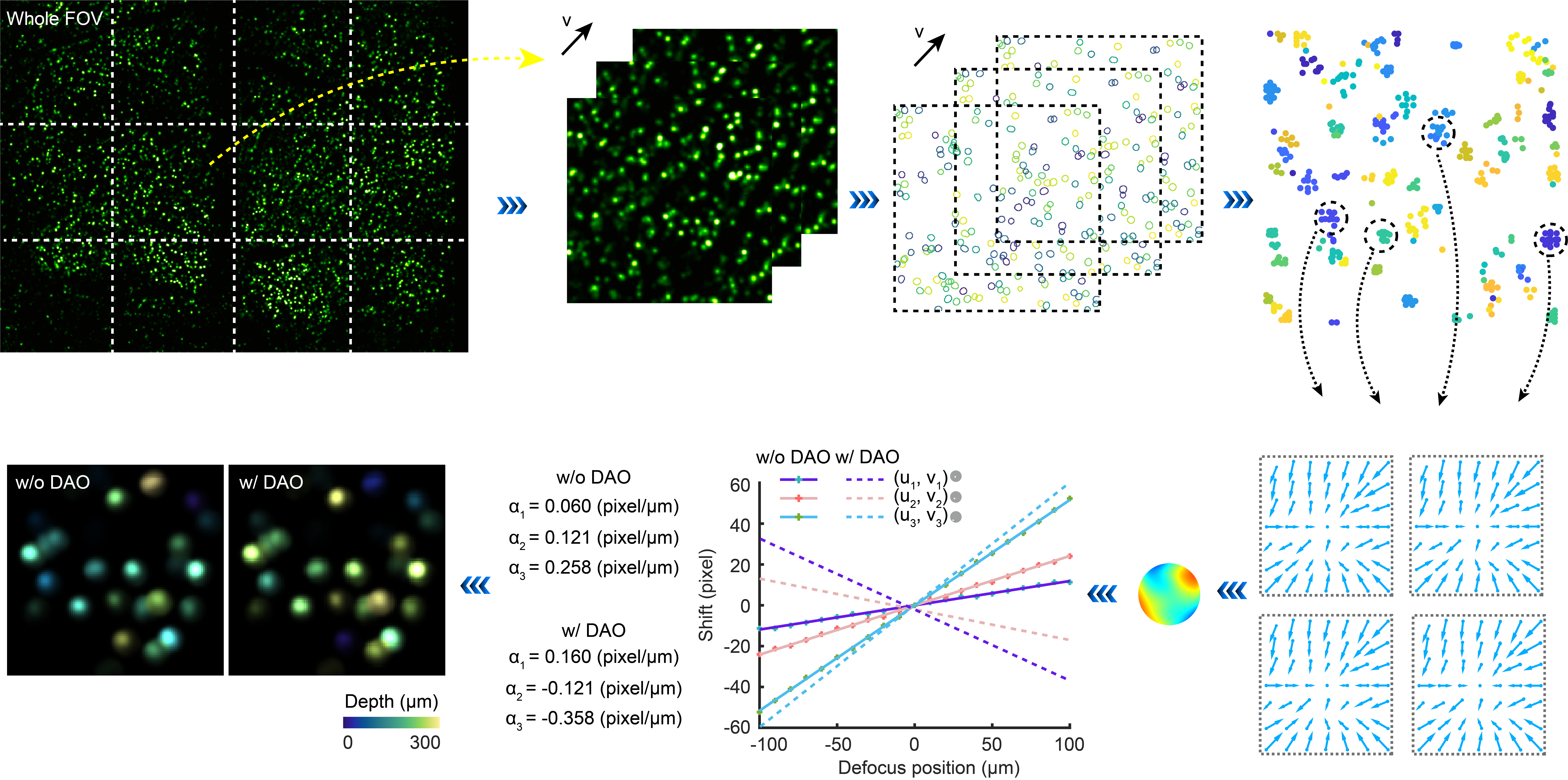

**Supplementary Figure 4 | Principle of aberration correction based on multi-view neuronal consistency.** To mitigate spatial localization errors caused by phase aberrations in light field recordings, the field of view is partitioned into a grid of 3×4 patches for each view. Within each patch, neuronal spatial footprints and their associated temporal calcium traces are extracted after background suppression using DeepWonder3D . For each neuron, apparent positional shifts across different views are analyzed to estimate the local phase aberration. The phase correction for a given patch is calculated by averaging the phase estimates derived from all neurons within that patch. Based on these averaged phase values, a refined linear relationship between axial defocus and inter-view disparity is established. This calibration improves depth estimation by adjusting the computed axial position of each extracted neuron, thereby enhancing the accuracy of three-dimensional localization.

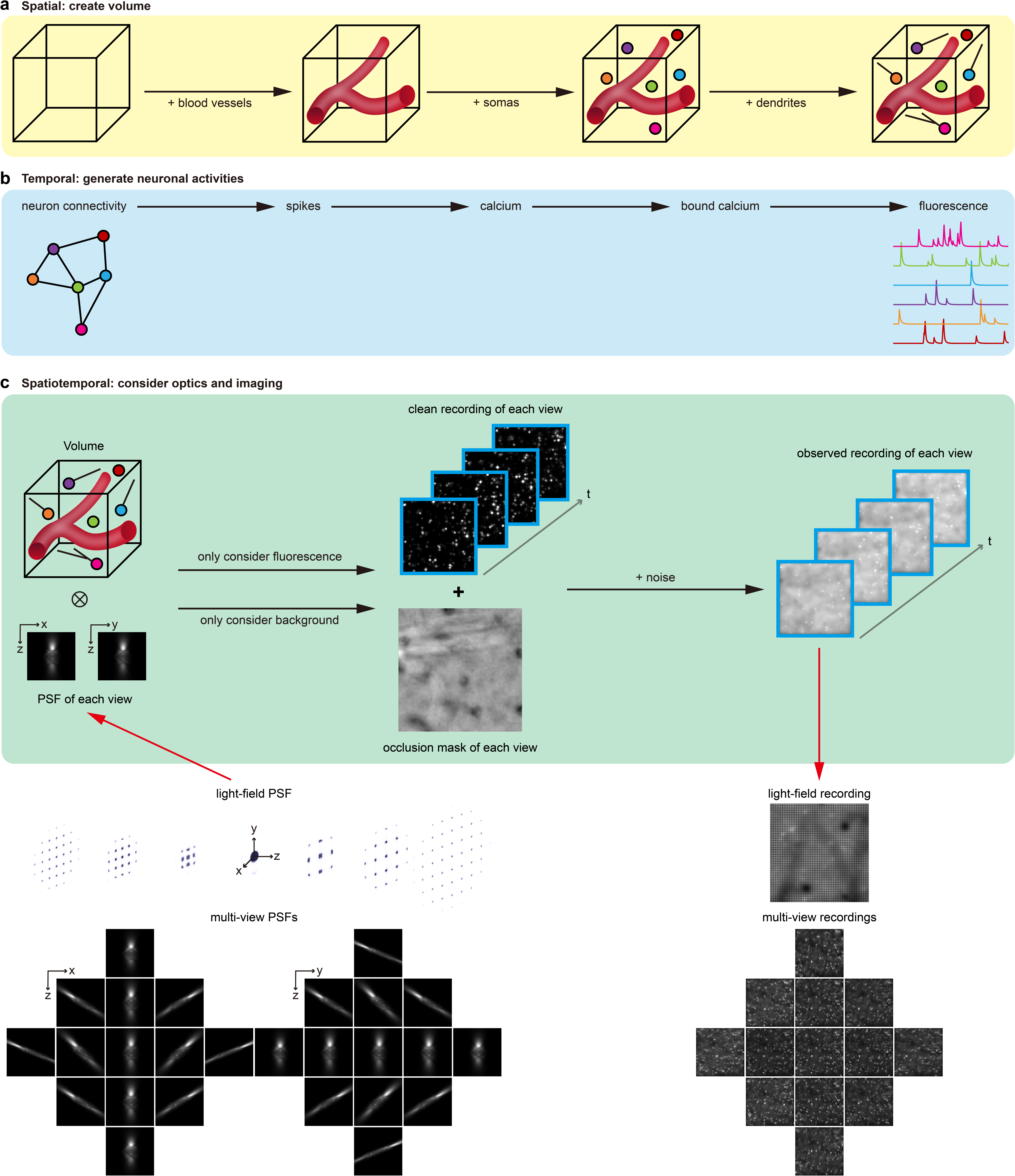

**Supplementary Figure 5 | NAOMi-LF workflow for generating simulated light field and multi-view calcium imaging datasets. a**, Construction of the spatial substrate. A computational 3D volume is initialized with static tissue structures, including blood vessels, neuronal somas, and dendrites. These components are sequentially embedded to prevent spatial overlap and collectively define the static spatial ground truth. **b**, Simulation of neuronal dynamics. A neuronal network model is used to generate synthetic spike trains. These spike events are convolved with models of calcium kinetics and fluorescence indicator dynamics, producing temporally resolved fluorescence signals for each neuron. **c**, Image formation modeling. The instantaneous 3D fluorescence volume (product of the spatial structure **a** and temporal activity **b**) is convolved with a view-specific point spread function (PSF) to simulate optical blurring, producing the clean image for each view. An occlusion mask is independently generated based on background light scattering. The simulated image frame results from a linear combination of the clean image, occlusion mask, and additive imaging noise. Iteration of this process over time yields final simulated recordings. By employing either a light field PSF or a set of multi-view PSFs, this workflow outputs corresponding simulated light field or multi-view calcium imaging recordings.

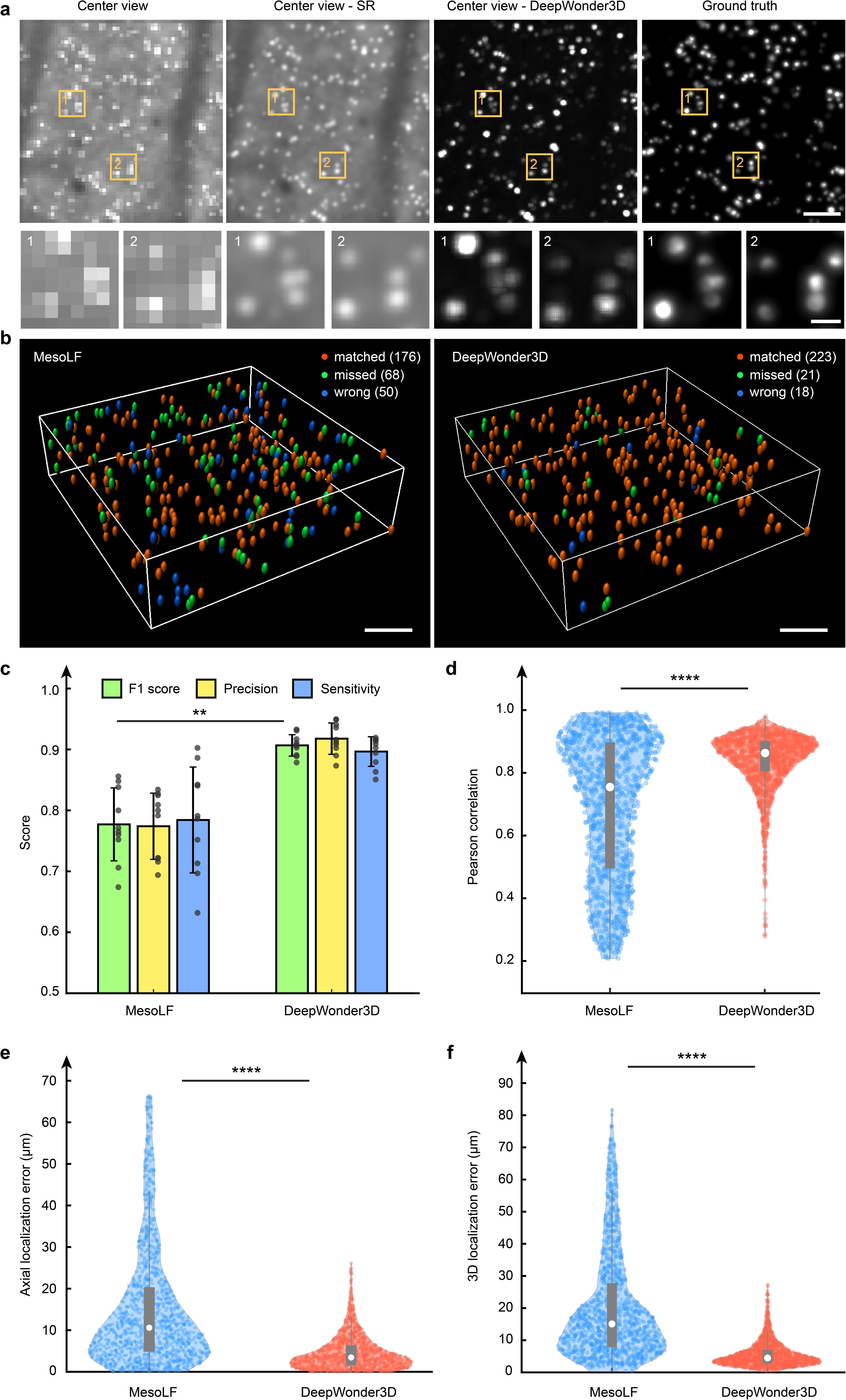

**Supplementary Figure 6 | Comparative evaluation of DeepWonder3D and MesoLF on simulated single-photon calcium imaging data. a**, Representative center-view images from simulated recordings, including raw input, super-resolved output, background-removed result from DeepWonder3D , and ground truth image (from left to right). Corresponding zoom-in panels are shown below each image. **b,** Comparison of neuronal extraction outcomes between MesoLF and DeepWonder3D . **c,** Quantitative assessment of extraction performance. DeepWonder3D achieves higher F1 score (0.91±0.02), precision (0.92±0.03), and sensitivity (0.90±0.02) than MesoLF (0.78±0.06, 0.77±0.05, 0.78±0.09, respectively). Data represent n = 10 independent simulated recordings. Bar height indicates the mean, error bars denote standard deviation (SD), and gray dots represent individual measurements. P = 0.002, two-sided Wilcoxon signed-rank test. **d.** Pearson correlation coefficients between extracted and ground-truth calcium traces. DeepWonder3D (orange, 0.84±0.09, n=1364 neurons from 7 simulated recordings) outperforms MesoLF (blue, 0.70±0.22, n=1181 neurons). **P < 0.0001, two-sided Wilcoxon rank-sum test. Visualized as violin plots with overlaid data points, median (white circle), interquartile range (thick gray bar), and proximal values (thin lines). **e.** Axial localization error across neurons. DeepWonder3D significantly reduces axial error (4.59 ± 4.14 μm) compared to MesoLF (15.19 ± 14.46 μm). Same statistical conventions as in **d**. **f.** 3D localization error comparison, with DeepWonder3D achieving substantially lower overall spatial error (5.61 ± 3.94 μm) relative to MesoLF (21.47 ± 16.73 μm). Same symbols and statistics as in **d**. Scale bars: 100 μm in panels **a** and **b**, 20 μm in zoom-in panels of **a**.

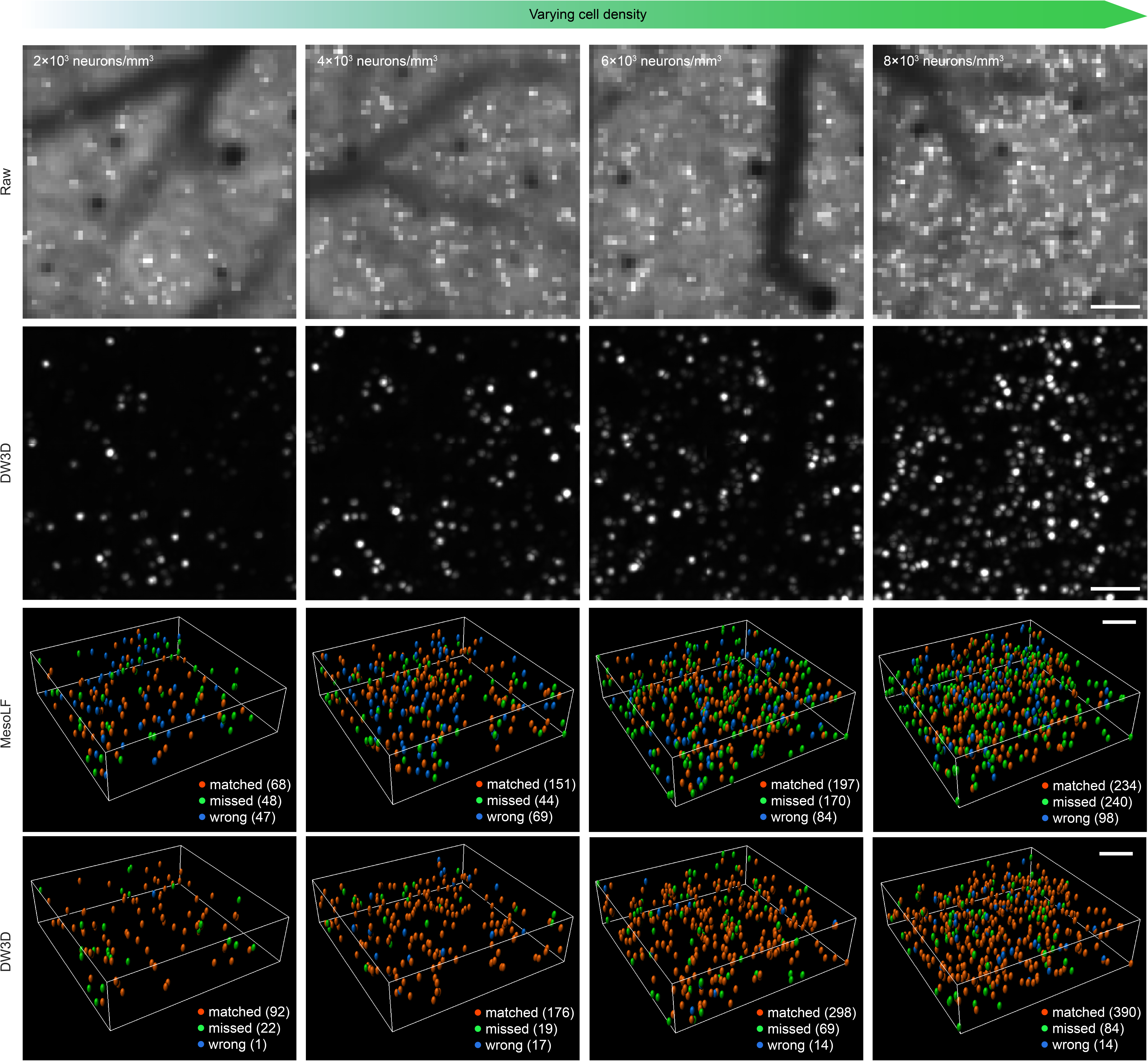

**Supplementary Figure 7 | Performance comparison between DeepWonder3D and MesoLF under varying neuronal densities.** Simulated multi-view recordings with increasing neuronal densities (2×10^3^，4×10^3^，6×10^3^，8×10^3^ neurons mm^-3^), generated using the NAOMi 1p simulator. First row: standard deviation projection over time of the center-view light field data. Second row: corresponding projections after background removal by DeepWonder3D . Third row: neuronal extraction results from MesoLF; orange dots indicate true positives (neurons correctly matched to ground truth), green dots indicate false negatives (missed neurons), and blue dots indicate false positives (erroneous extractions). Fourth row: extraction results from DeepWonder3D with identical color coding. Scale bars: 100 μm.

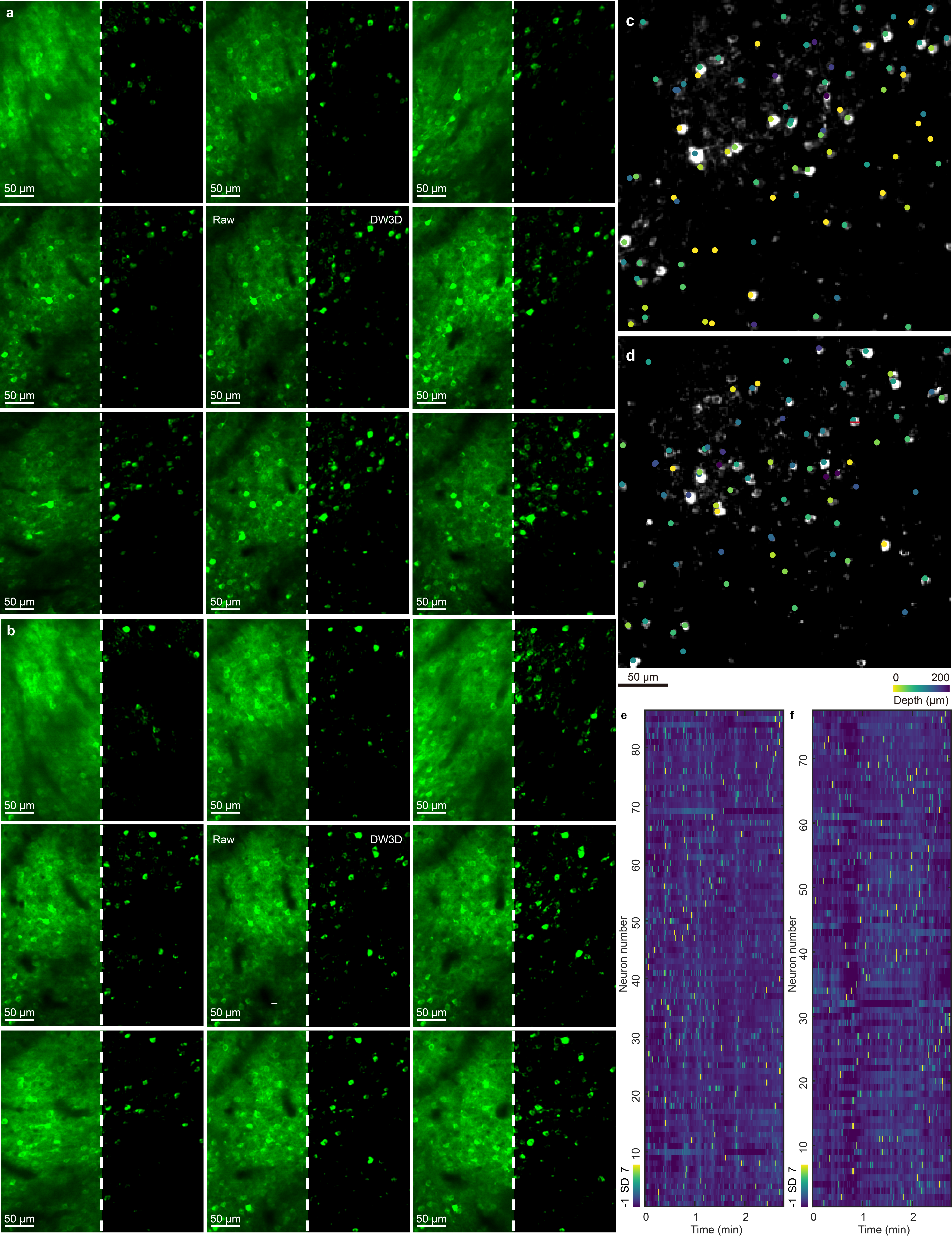

**Supplementary Figure 8 | Application of DeepWonder3D to multi-view two-photon *in vivo* calcium imaging data.** Data and results from two mice are presented: Mouse A (**a, c, e**) and Mouse B (**b, d, f**). a, b, Temporal standard deviation (STD) projections from multiple views before (left panels) and after (right panels) DeepWonder3D processing. c, d, 3D reconstruction of neuron positions from data in (**a, b**). Depth-encoded centroids of all extracted neurons (colored points) are overlaid on the temporal-STD projection of central view. e, f, Heatmaps of ΔF/F traces for all neurons extracted by DeepWonder3D over the two 167-second recordings. Each row represents the activity of a single neuron. Scale bars: 50 μm (**a, b, c, d**).

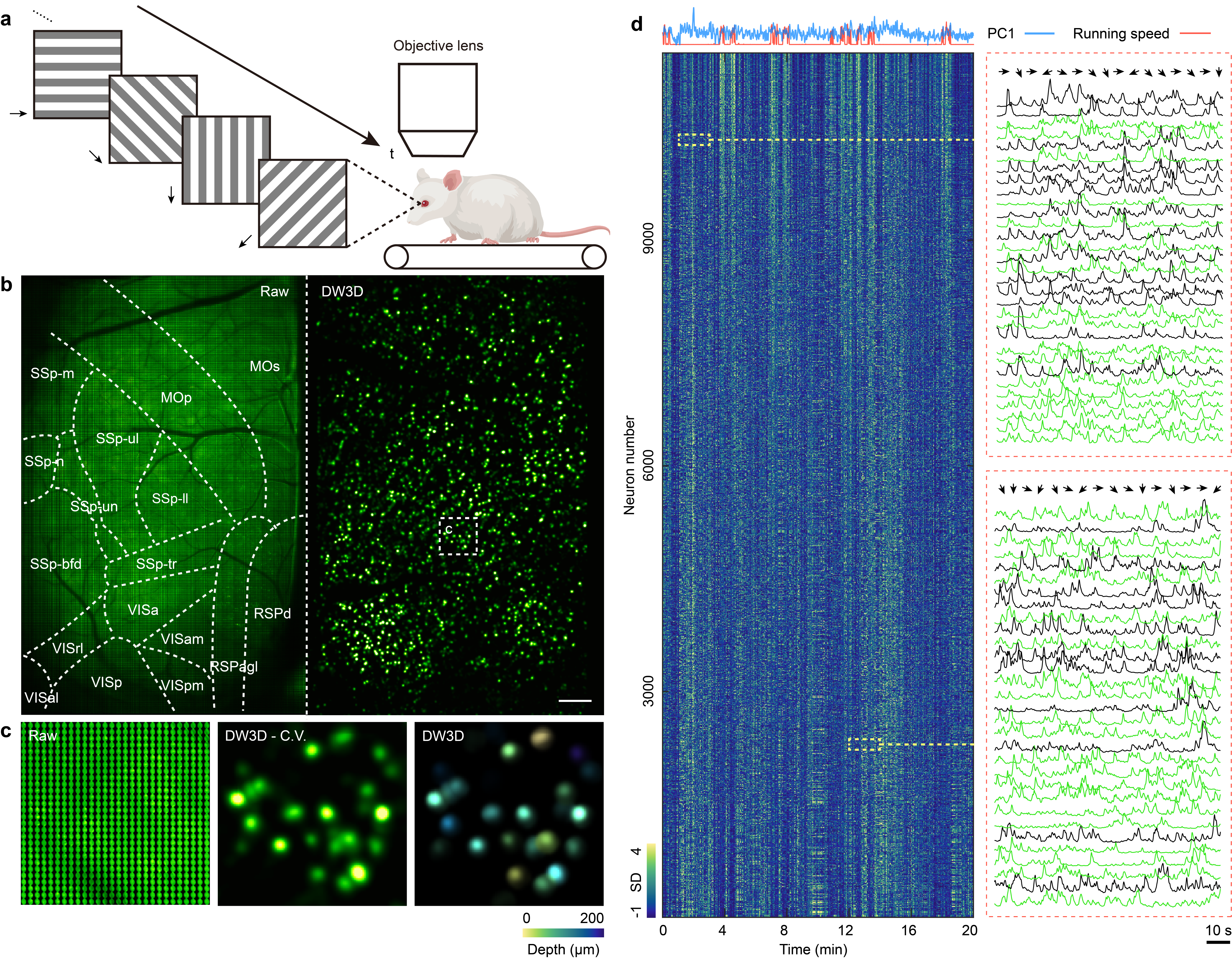

**Supplementary Figure 9 | Application of DeepWonder3D to large-scale *in vivo* optical neurophysiology data. a**, Schematic of the experimental setup. A head-fixed mouse was presented with full-field drifting gratings oriented in eight directions (22.5° increments over 180°). Visual stimuli are illustrated as a sequence of gratings with arrows indicating motion direction. Simultaneously, neuronal calcium activity across the dorsal cortex was recorded using a light field microscope. **b**, Widefield calcium imaging results. Left: center-view projection of raw light field data, overlaid with cortical region boundaries. Right: corresponding result after background removal using DeepWonder3D . **c**, Zoom-in view of the white dashed box in **b**, showing the raw image, background-removed image, and depth-coded neuronal activity map reconstructed by DeepWonder3D . **d**, Heatmap of ΔF/F traces extracted from over 10,000 neurons during a 20-minute recording. Top traces represent the animal’s running speed (orange) and the first principal component (PC1) of neuronal activity (blue). Two enlarged panels highlight example calcium activity from 30 neurons each. Arrows denote epochs corresponding to different grating orientations. Scale bars: 400 μm in panel **b**, 10 s in panel **d**.

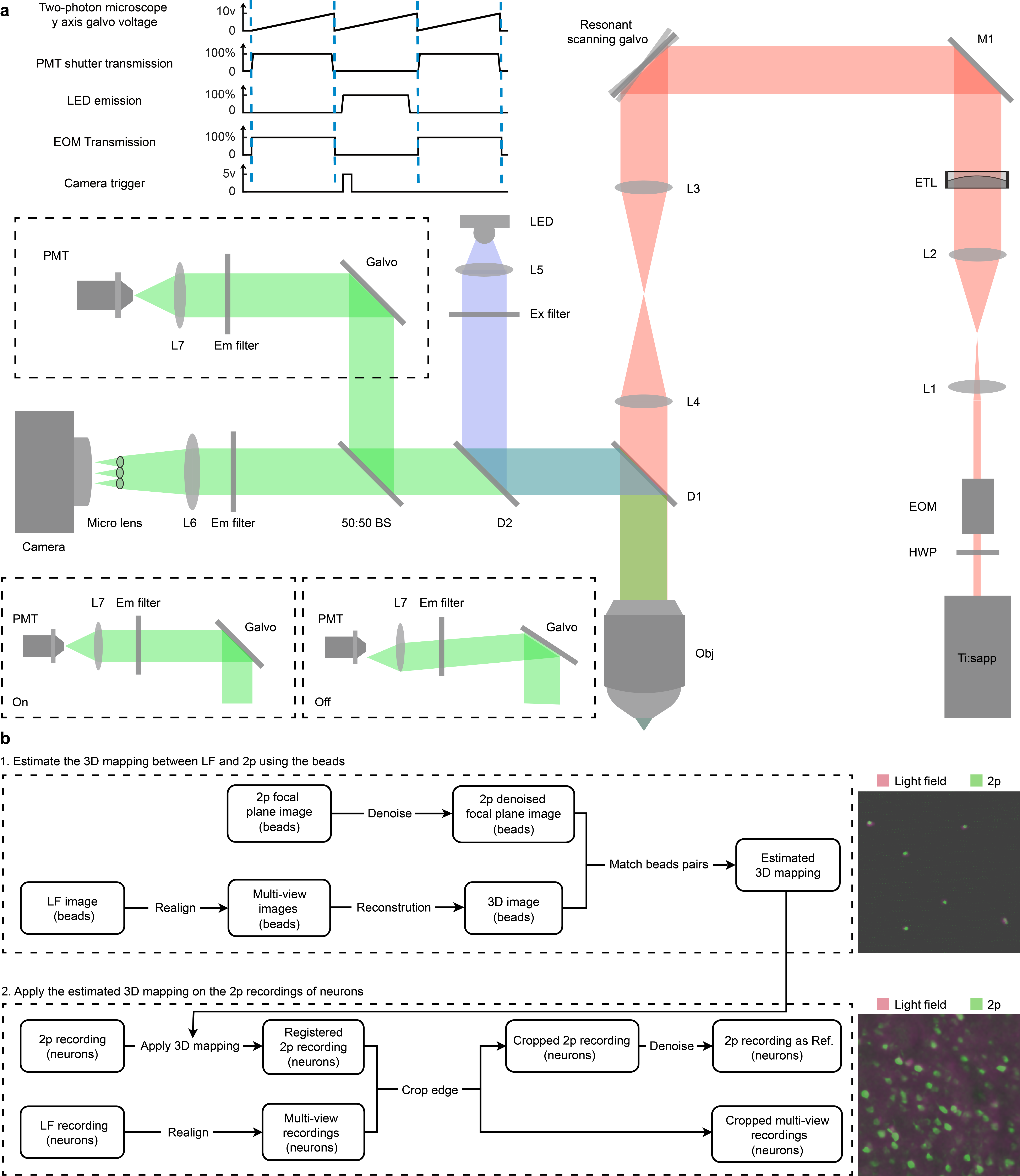

**Supplementary Figure 10 | Schematic of the hybrid light field and two-photon microscope for simultaneous neuronal activity recording. a**, Optical layout of the custom-built dual-modality imaging system integrating a traditional light field microscope with a two-photon excitation path. Ti:sapp: tunable titanium-sapphire laser source; HWP: half-wave plate; EOM: electro-optic modulator; ETL: electrically tunable lens; M: mirror; L1-L7: lenses; D1: long-pass dichroic mirror separating emission fluorescence (520 nm, green path) and light field excitation light (470 nm, blue path) from two-photon excitation light (920 nm, orange path); D2: long-pass dichroic mirror separating fluorescence signals from the light field excitation path. BS: 50:50 non-polarizing beam splitter; Em filter: 520 nm emission filter; Ex filter: 475 nm excitation filter; PMT: photomultiplier tube. Bottom left inset: The galvo mirror acts as an optical shutter, enabling dynamic control of the light path to the PMT; both "on" and "off" states are illustrated. Top left inset: Timing diagram for synchronizing key components, including the y-axis galvo for two-photon scanning, the galvo-based shutter before the PMT, LED excitation, EOM modulation, and the camera trigger signal. **b**, Field-of-view (FOV) co-registration workflow for the light field and two-photon channels. Fluorescent bead images are acquired simultaneously at the two-photon focal plane to establish spatial correspondence. An affine transformation is derived from manually annotated feature point pairs across modalities, enabling accurate lateral mapping of the two-photon FOV onto the light field FOV. This alignment allows the high-resolution two-photon calcium imaging data to serve as a ground-truth reference for validating neuronal extraction and activity extraction in the light field recordings.

### References

1.	NoRMCorre: An online algorithm for piecewise rigid motion correction of calcium imaging data. *J. Neurosci. Methods* **291,** 83–94 (2017).

2.	Giovannucci, A., Friedrich, J., Gunn, P., Kalfon, J., Brown, B. L., Koay, S. A., Taxidis, J., Najafi, F., Gauthier, J. L., Zhou, P., Khakh, B. S., Tank, D. W., Chklovskii, D. B. & Pnevmatikakis, E. A. CaImAn an open source tool for scalable calcium imaging data analysis. *eLife* **8,** 1–45 (2019).
